# Genome-wide DNA methylation and gene expression patterns reflect genetic ancestry and environmental differences across the Indonesian archipelago

**DOI:** 10.1101/704304

**Authors:** Heini Natri, Katalina S. Bobowik, Pradiptajati Kusuma, Chelzie Crenna Darusallam, Guy S. Jacobs, Georgi Hudjashov, J. Stephen Lansing, Herawati Sudoyo, Nicholas E. Banovich, Murray P. Cox, Irene Gallego Romero

**Affiliations:** Center for Evolution and Medicine, School of Life Sciences, Arizona State University, Tempe 85281, AZ, USA; The Translational Genomics Research Institute, Phoenix 85004, AZ, USA; Melbourne Integrative Genomics, University of Melbourne, Parkville 3010, Australia; School of BioSciences, University of Melbourne, Parkville 3010, Australia; Centre for Stem Cell Systems, University of Melbourne, Parkville 3010, Australia; Genome Diversity and Diseases Laboratory, Eijkman Institute for Molecular Biology, Jakarta 10430, Indonesia; Complexity Institute, Nanyang Technological University, Singapore 637723, Singapore; Statistics and Bioinformatics Group, School of Fundamental Sciences, Massey University, Palmerston North 4410, New Zealand; Santa Fe Institute, Santa Fe, NM 87501, USA; Vienna Complexity Science Hub, Vienna 1080, Austria; Stockholm Resilience Center, Kräftriket, Stockholm 10405, Sweden; Department of Medical Biology, Faculty of Medicine, University of Indonesia, Jakarta 10430, Indonesia; Sydney Medical School, University of Sydney, Sydney, NSW 2006, Australia

**Keywords:** Indonesia, RNA-sequencing, DNA methylation, gene expression, molecular phenotypes

## Abstract

Indonesia is the world’s fourth most populous country, host to striking levels of human diversity, regional patterns of admixture, and varying degrees of introgression from both Neanderthals and Denisovans. However, it has been largely excluded from the human genomics sequencing boom of the last decade. To serve as a benchmark dataset of molecular phenotypes across the region, we generated genome-wide CpG methylation and gene expression measurements in over 100 individuals from three locations that capture the major genomic and geographical axes of diversity across the Indonesian archipelago. Investigating between- and within-island differences, we find up to 10% of tested genes are differentially expressed between the islands of Mentawai (Sumatra) and New Guinea. Variation in gene expression is closely associated with DNA methylation, with expression levels of 9.7% of genes strongly correlating with nearby CpG methylation, and many of these genes being differentially expressed between islands. Genes identified in our differential expression and methylation analyses are enriched in pathways involved in immunity, highlighting Indonesia tropical role as a source of infectious disease diversity and the strong selective pressures these diseases have exerted on humans. Finally, we identify robust within-island variation in DNA methylation and gene expression, likely driven by very local environmental differences across sampling sites. Together, these results strongly suggest complex relationships between DNA methylation, transcription, archaic hominin introgression and immunity, all jointly shaped by the environment. This has implications for the application of genomic medicine, both in critically understudied Indonesia and globally, and will allow a better understanding of the interacting roles of genomic and environmental factors shaping molecular and complex phenotypes.

## Introduction

Modern human genomics does not equitably represent the full breadth of humanity. While genome sequences for people of European descent now number a million or more, most of the world is deeply understudied^1^. This is particularly true of Indonesia^2^, a country geographically as large as continental Europe and the world’s fourth largest by population. Genomic diversity in Indonesia is strikingly different to other well-characterized East Asian populations, such as Han Chinese and Japanese, but this diversity is not captured in large global datasets like the 1000 Genomes Project^3^ or the Simons Genome Diversity Project^4^. The first Indonesian genome sequences were only reported in 2016^5^ with the first representative survey of diversity across the archipelago only appearing in 2019^6^. This extreme lack of representation extends to molecular phenotypes. To our knowledge, only one genome-wide gene expression study has been published^7^ from the region, focused exclusively on host-pathogen interactions with *P. falciparum.* There are no analyses of diversity in gene regulatory mechanisms in either Indonesia or, more broadly, Island Southeast Asia.

This gap is especially incongruous because Indonesia is an epicenter of infectious disease diversity, ranging from well-known agents like malaria^8^ to emerging diseases like zika virus^9^. The country faces substantial healthcare challenges, including the rise in prevalence of understudied tropical infectious diseases and the increasing impact of metabolic disorders among a growing middle class^10^. However, Indonesia also offers unique advantages for studying responses to these diseases and disorders, some of which are likely to have exerted strong evolutionary pressures on the immune system over thousands of years^11^. Because the country comprises a chain of islands that stretch for 50 degrees of longitude along the equator (wider than either the continental USA or mainland Europe), but span barely 15 degrees of latitude, environment conditions are broadly comparable in many key respects across Indonesia. In contrast, a complex population history means that its people differ greatly, forming a genomic cline from Asian ancestry in the west to Papuan ancestry in the east^12^. This change in ancestry is the most distinctive genomic signal observed in the region^13^, and provides a framework for studying the effects of genome composition on gene expression in a heterogeneous environment.

To provide a benchmark dataset of regional molecular phenotypes, here we report genome-wide measurements of DNA methylation and gene expression for 117 individuals drawn from three population groups that capture the major genomic and geographical axes of diversity across Indonesia. The people of Mentawai, living on the barrier islands off Sumatra, are representative of the dominant Asian ancestry in western Indonesia^13^; the Korowai, hunter-gatherers from the highlands of western New Guinea capture key aspects of regional Papuan ancestry^6^; and the inhabitants of Sumba in eastern Indonesia are, genetically, a near equal mixture of the two different ancestries^14^. However, it remains unclear whether, and to what extent, these differences in genomic ancestry correlate with variation in molecular phenotypes. By quantifying DNA methylation and gene expression levels across Indonesia for the first time, we identify the relative influences of genomic ancestry versus plasticity to local environmental conditions in driving regional molecular phenotypic patterns.

## Methods

### Ethical approvals

The samples used in this study were collected by JSL, HS and an Indonesian team from the Eijkman Institute for Molecular Biology, Jakarta, Indonesia, with the assistance of Indonesian Public Health clinic staff. All collections followed protocols for the protection of human subjects established by institutional review boards at the Eijkman Institute (EIREC #90 and EIREC #126) and the University of Melbourne (Human Ethics Sub-Committee approval 1851639.1). All individuals gave written informed consent for participation in the study. Permission to conduct research in Indonesia was granted by the Indonesian Institute of Sciences and by the Ministry for Research, Technology and Higher Education.

### Data collection

Whole blood was collected by trained phlebotomists from the Eijkman Institute from over 300 Indonesian men. Samples were collected across multiple villages in the three islands using EDTA blood tubes from either Vacuette or Intherma for DNA isolation, and Tempus Blood RNA Tubes (Applied Biosystems) for RNA isolation. All RNA extractions were performed according to the manufacturers’ protocols and randomised with respect to village and island (Supplementary Tables 1 and 2).

Quality and concentration of all extracted RNA samples were assessed with a Bioanalyzer 2100 (Agilent) and a Qubit device (Life Technologies), respectively. We selected 117 samples for RNA sequencing and DNA methylation analysis primarily on the basis of RIN score, by focusing on villages with at least 10 samples with RIN ≥ 6 (Table 1). Given our past work on the island of Sumba^14^, we included all samples from Sumba with RIN ≥ 6, heedless of village. However, we occasionally observed differences between our RIN measurements and those performed by the sequencing provider, with the latter generally being lower. Out of 117 individuals, 24 (21%) had a final RIN measurement < 6. Further detail on all samples, including extracting and sequencing batches, is provided in Supplementary Tables 1 and 2. Library preparation was performed by Macrogen (South Korea), using 750 ng of RNA and the Globin-Zero Gold rRNA Removal Kit (Illumina) according to the manufacturer’s instructions. Samples were sequenced using a 100-bp paired-end configuration on an Illumina HiSeq 2500 to an average depth of 30 million read pairs per individual, in three batches. All batches included at least one inter-batch control for downstream normalisation (Supplementary Tables 1 and 2).

**Table 1:**
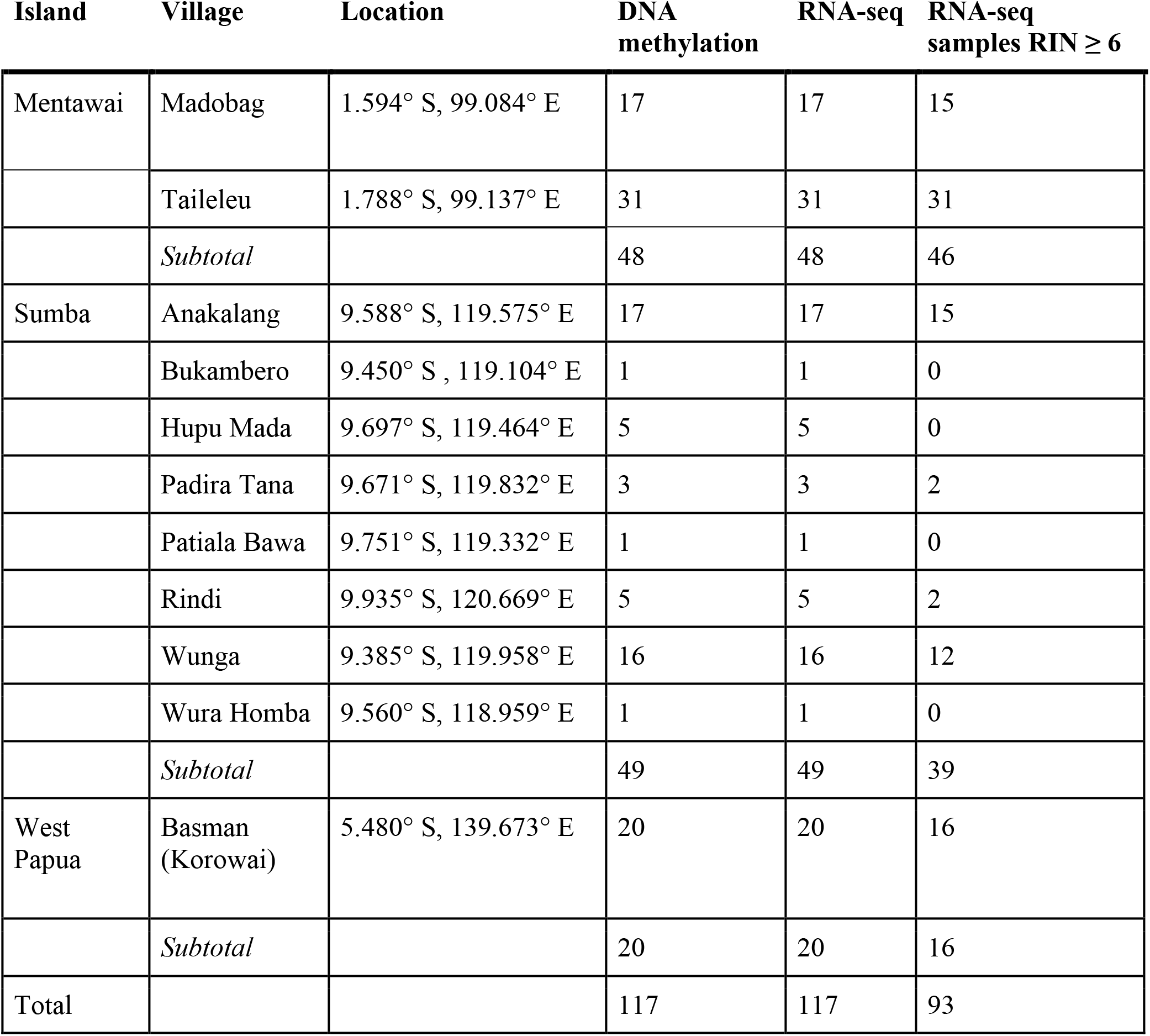
Numbers of DNA methylation and RNA sequenced samples from each study location.

| Island | Village | Location | DNA<br>methylation | RNA-seq | RNA-seq<br>samples RIN ≥ 6 |
| --- | --- | --- | --- | --- | --- |
| Mentawai | Madobag | 1.594° S, 99.084° E | 17 | 17 | 15 |
|  | Taileleu | 1.788° S, 99.137° E | 31 | 31 | 31 |
|  | *Subtotal* |  | 48 | 48 | 46 |
| Sumba | Anakalang | 9.588° S, 119.575° E | 17 | 17 | 15 |
|  | Bukambero | 9.450° S, 119.104° E | 1 | 1 | 0 |
|  | Hupu Mada | 9.697° S, 119.464° E | 5 | 5 | 0 |
|  | Padira Tana | 9.671° S, 119.832° E | 3 | 3 | 2 |
|  | Patiala Bawa | 9.751° S, 119.332° E | 1 | 1 | 0 |
|  | Rindi | 9.935° S, 120.669° E | 5 | 5 | 2 |
|  | Wunga | 9.385° S, 119.958° E | 16 | 16 | 12 |
|  | Wura Homba | 9.560° S, 118.959° E | 1 | 1 | 0 |
|  | *Subtotal* |  | 49 | 49 | 39 |
| West<br>Papua | Basman<br>(Korowai) | 5.480° S, 139.673° E | 20 | 20 | 16 |
|  | *Subtotal* |  | 20 | 20 | 16 |
| Total |  |  | 117 | 117 | 93 |

In parallel, we extracted whole blood DNA from all individuals included in the RNA sequencing data using Gentra^®^ Puregene^®^ for human whole blood kit (QIAGEN) and MagAttract^®^ HMW DNA kit (QIAGEN) according to the manufacturer’s instructions. 1 μg of DNA from each sample was shipped to Macrogen, bisulfite-converted and hybridized to Illumina Infinium EPIC BeadChips according to the manufacturer’s instructions. Samples were randomized with respect to village and island across two array batches, with three samples processed on both batches to control for technical variation (Supplementary Table 1).

### RNA sequencing data processing

All RNA sequencing reads were examined with FastQC v. 0.11.5^15^. Leading and trailing bases below a Phred score of 20 were removed using Trimmomatic v. 0.36^16^. Reads were then aligned to the human genome (GRCh38 Ensembl release 90: August 2017) with STAR v. 2.5.3a^17^ and a two-pass alignment mode; this resulted in a mean of ~29 million uniquely-mapped read pairs per sample. Next, we performed read quantification with featureCounts v. 1.5.3^18^ against a subset of GENCODE basic (release 27) annotations that included only transcripts with support levels 1-3, retaining a total of 58,391 transcripts across 29,614 genes. On average, we successfully assigned ~15 million read pairs to each sample (Supplementary table 2).

### Differential expression analysis

All statistical analyses were performed using R v. 3.5.2^19^. We transformed read counts to log2-counts per million (CPM) using a prior count of 0.25, and removed genes with low expression levels by only keeping genes with log2 CPM ≥ 1 in at least half of the individuals from any island, resulting in a total of 12,975 genes retained for further analysis. To quantify the effect of technical batch, we included six replicate samples among our sequencing batches. As expected, PCA of uncorrected data suggested the presence of substantial sequencing batch effects in the data (Supplementary figure 1). However, pairwise correlations between technical replicates were higher than between different individuals from the same village sequenced in the same batch (Supplementary figure 2).

We applied TMM normalisation^20^ to the data, and removed high sample variability from the count data using the *voom* function^21^ in limma v. 3.40.2^22^. Differential expression testing was also performed using limma. To construct the linear model for testing, we used ANOVA to test for associations between all possible covariates and the first 10 principal components (PC) of the data. Technical covariates significantly associated with at least one PC (sequencing batch, RIN, age) were included in the model. In addition, because blood cell type composition can impact gene expression estimates in bulk RNA samples, we used DeconCell v. 0.1.0^23^ to estimate the proportion of CD8T, CD4T, NK, B cells, monocytes and granulocytes in each sample (Supplementary table 2), and tested these for association with the first 10 PCs as described above. All covariates were significantly associated with at least one PC and were included in the differential expression model. Sampling sites were included at either the island or the village level, depending on the test. Comparisons between villages were limited to those with at least 15 individuals, to ensure sufficient power to detect differences. All individuals were included in comparisons between islands, and models were not hierarchically structured. Genes were called as differentially expressed (DEG) if the FDR-adjusted *p* value was below 0.01, regardless of the magnitude of the log2 fold change, unless noted otherwise.

Lists of DEGs were annotated using biomaRt v. 2.40.0^24^. Gene set enrichment analyses for the DEGs on the island and village levels were performed using clusterProfiler v. 3.12.0^25^, with Gene Ontology and KEGG annotation drawn from the org.Hs.eg.db v. 3.9 database^26^. Additionally, we tested whether DEGs were enriched for genes known to have been introgressed from Denisovans into individuals of Papuan ancestry at high frequency using a hypergeometric test. A GO term similarity test was performed using GOSim v. 1.22.0^27^ using the ‘relevance’ method. Finally, to examine possible associations between known climatic variables and expression across sampling sites, we retrieved mean monthly precipitation and temperature data from WorldClim v. 2.0^28^ for the five main villages in our study at a resolution of 0.5 arcminutes (roughly 1 km^2^ tiles).

### DNA methylation array data processing and analysis

DNA methylation data were processed using minfi v. 1.30.0^29^. The two arrays were combined using the *combineArrays* function and preprocessed with the *bgcorrect.illumina* function to correct for array background signal. Signal strength across all probes was evaluated using the *detectionP* function and probes with signal *p* < 0.01 in >75% of samples were retained. To avoid potential spurious signals due to differences in probe hybridization affinity, we discarded 6,072 probes overlapping known SNPs segregating in any of the study populations based on previously published genotype data^6^. The final number of probes retained was 859,404. Subset-quantile Within Array Normalization (SWAN) was carried out using the *preprocessSWAN* function^30^. Methylated and unmethylated signals were quantile normalized using lumi v. 2.36.0^31^. As with the RNA sequencing, replicate samples were included to detect and correct for batch effects (supplementary figure 3). The replicate samples exhibit a high correlation between batches (Spearman’s Rho 0.969 for MPI-025 and 0.980 for SMB-ANK-029, Supplementary Figure 4). As above, we used limma to test for differential methylation between sampling sites. We included methylation array batch, age, and the estimated cell type proportions (derived from the RNA sequencing data) as covariates. Differentially methylated probes (DMPs) between all pairwise comparisons of the islands and villages were identified using contrast designs. Significant DMPs were selected based on an FDR-adjusted *p* value threshold of 0.01 and a log2 fold change of 0.5 or greater. Enrichment tests for the DMPs were performed using missMethyl v. 1.18.0^32^, to account for differences in probe density associated with gene length that can otherwise bias results^33^; probes were annotated to genes according to Illumina’s manifest for the EPIC array.

We further identified differentially methylated regions (DMRs) by annotating the CpG probes with the *cpg.annotate* function of the R package DMRcate v. 3.9^34^, and by collapsing the probes to regions using the *dmrcate* function. Individual probes with an FDR-adjusted *p* value ≤0.01 and significant DMRs were selected based on a region beta value of 0.5 or greater.

### Principal Component Analysis (PCA)

DNA methylation M-values and gene expression log_2_ CPM values were adjusted to correct for batch effects and differences in blood cell type proportions between samples by fitting a linear model with the technical covariates used in the differential methylation and expression analysis. Residuals of this model were used in the PCAs in Figure 1. Variable CpG probes and genes were identified based on coefficients of variation between samples. PCA was performed using the 10^4^ most variable probes and the 10^3^ most variable genes from the methylation and expression datasets, respectively; PCAs of the entire data set before and after batch correction are available in supplementary figures 1 and 3.

**Figure 1.**
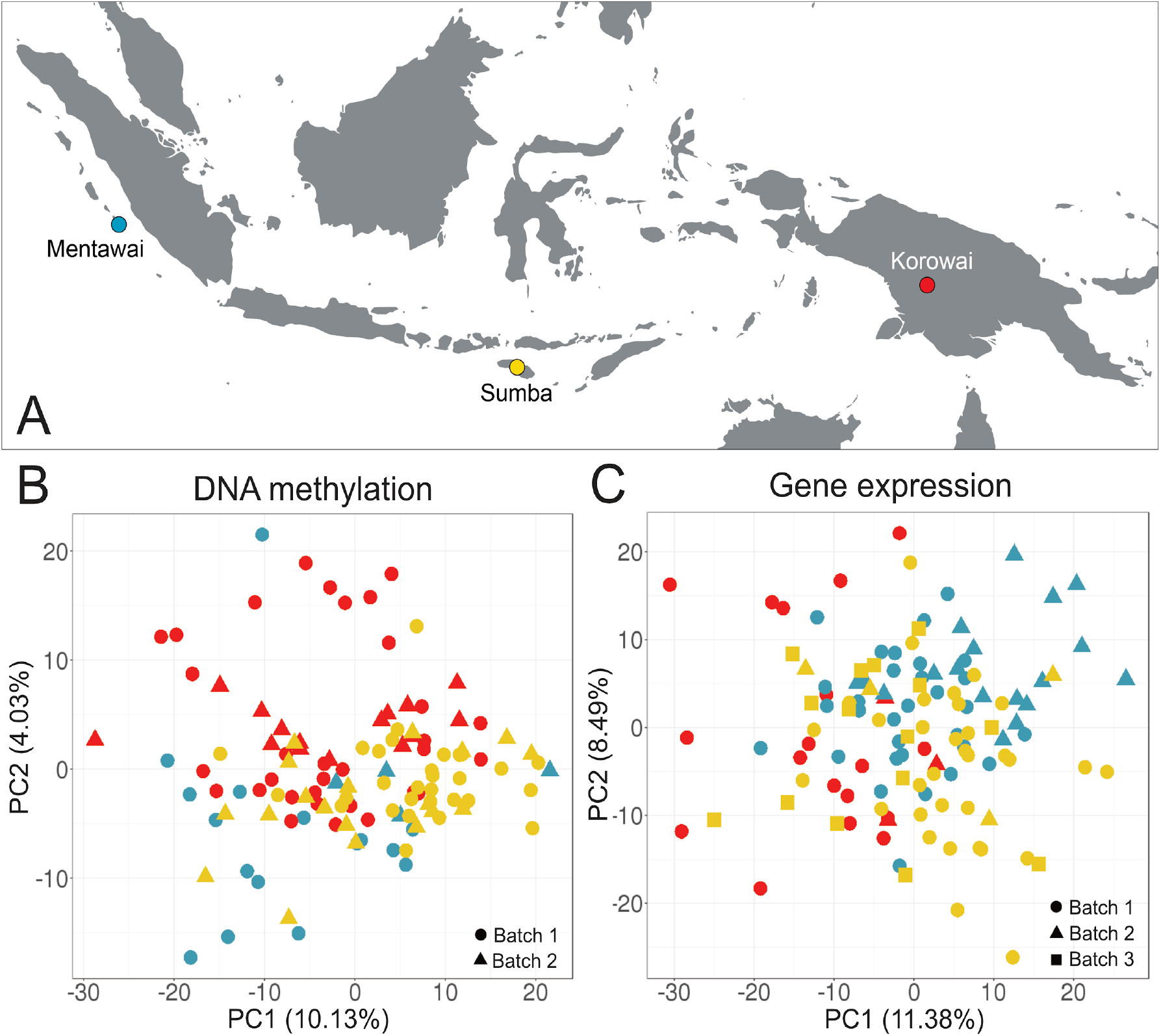
Sampling locations and overview of DNA methylation and gene expression variation among the study samples. (A) Colors indicate island populations: Mentawai, blue; Sumba, yellow; Korowai, red. PCA was performed on the top 10,000 most variable methylation probes and the top 1,000 most variable genes, determined by the sample-wide coefficient of variation. The first two axes of variation from the principal component analysis in the (B) DNA methylation and (C) gene expression data after correcting for confounding effects are driven by between-island differences. Plotting shapes indicates sequencing/array batches.

### Identifying associations between DNA methylation regions and gene expression

We used the R package MethylMix v. 2.12.0^35,36^ to identify transcriptionally predictive methylation states by focusing on methylation changes that affect gene expression. As with the PCA analysis, DNA methylation M-values and gene expression log (CPM) values were adjusted to account for technical covariates and blood cell type proportions by fitting a linear model. Residuals of these linear models were used in the analysis. Batch corrected M-values and logCPM values were min-max normalized to range from 0 to 1. CpG probe methylation levels were matched to genes using the *ClusterProbes* function, which uses a complete linkage hierarchical clustering algorithm for all probes of a single gene to cluster the probes. To identify transcriptionally predictive DNA methylation events, MethylMix utilizes linear regression to detect negative correlations between methylation and gene expression levels. Matching DNA methylation and gene expression data from 117 individuals were used in the analysis, and a total of 10,420 genes with matching methylation and expression data were tested. As MethylMix does not output detailed summary statistics of the fitted linear models, we used linear regression to calculate the r^2^ and *p* values for each significant CpG probe cluster and gene pair detected by MethylMix. False discovery rate adjusted *p* values were calculated using the *p. adjust* function in base R.

### Data access

All RNA sequencing reads and Illumina Epic iDat files are available through the Data Access Committee of the official data repository at the European Genome-phenome Archive (EGA; https://www.ebi.ac.uk/ega/home). The RNA sequencing data are deposited in study EGAS00001003671 and the methylation data are deposited in study EGAS00001003653. Matrices of unfiltered read counts (doi:10.26188/5d12023f77da8) and M-values (doi:10.26188/5d13fb401e305) for all samples, including replicates, are freely available on figshare (https://figshare.com). Differential expression (doi:10.26188/5d26aec1d817a) and methylation (10.26188/5d26b0b5230dd) testing results are freely available on figshare.

## Results

### Differential DNA methylation and gene expression between Indonesian island populations

To quantify the gene regulatory landscape in Indonesia, we generated DNA methylation (array) and gene expression (RNA sequencing) measurements from 117 whole blood samples of male individuals living on three islands in the Indonesian archipelago (Figure 1A). Our three sampling sites, Mentawai, Sumba, and West Papua, represent distinct points along a well-documented Asian/Papuan admixture cline^13^: the Korowai of West Papua exhibit high Papuan ancestry; Sumbanese have intermediate degrees of Papuan ancestry; and the Mentawai have no Papuan ancestry, having been settled primarily by ancestral Austronesian speakers. Furthermore, Korowai individuals are likely to carry up to 5% of introgressed genomic sequence from archaic Denisovans, as repeatedly observed in other samples from the island of New Guinea^6,37^.

Principal component analysis of DNA methylation (Figure 1B) and gene expression (Figure 1C) shows clear clustering of samples driven by population origin. After correcting for known technical confounders, PC1 in the DNA methylation data separates the island of Sumba from both the Korowai (FDR-corrected ANOVA *p* = 0.001) and Mentawai (*p* = 7.4×10^−5^); PC2 further differentiates Sumbanese and Mentawai (*p* = 9.0×10^−4^) and additionally separates Mentawai from Korowai (*p* = 9.0×10^−7^). In the gene expression data, Korowai is separated from both Mentawai and Sumba (*p* = 1.0×10^−7^ and 1.5×10^−6^, respectively), whereas PC2 separates Sumba from Mentawai (*p* = 1.6×10^−4^).

We then tested for differences in DNA methylation and gene expression between the three islands, initially without considering the village structure in Sumba and Mentawai (Table 1; supplementary tables 1 and 2). At an absolute log2(FC) threshold of 0.5 and an FDR-adjusted *p* value threshold of 0.01, we detected 22,189 (2.58% of all tested probes), 14,168 (1.64%) and 3,947 (0.46%) differentially methylated probes (DMPs) and 1,398 (10.77% of all tested genes), 1,017 (7.84%), and 314 (2.40%) differentially expressed genes (DEGs) between Sumba and the Korowai, Mentawai and the Korowai, and Sumba and Mentawai, respectively (Figure 2A, 2B). In addition, we identified 1,003, 919 and 283 differentially methylated regions across all three inter-island comparisons, respectively, when thresholding to a mean β difference of 0.05 across the region. A full summary of these results is available as supplementary table 3.

**Figure 2.**
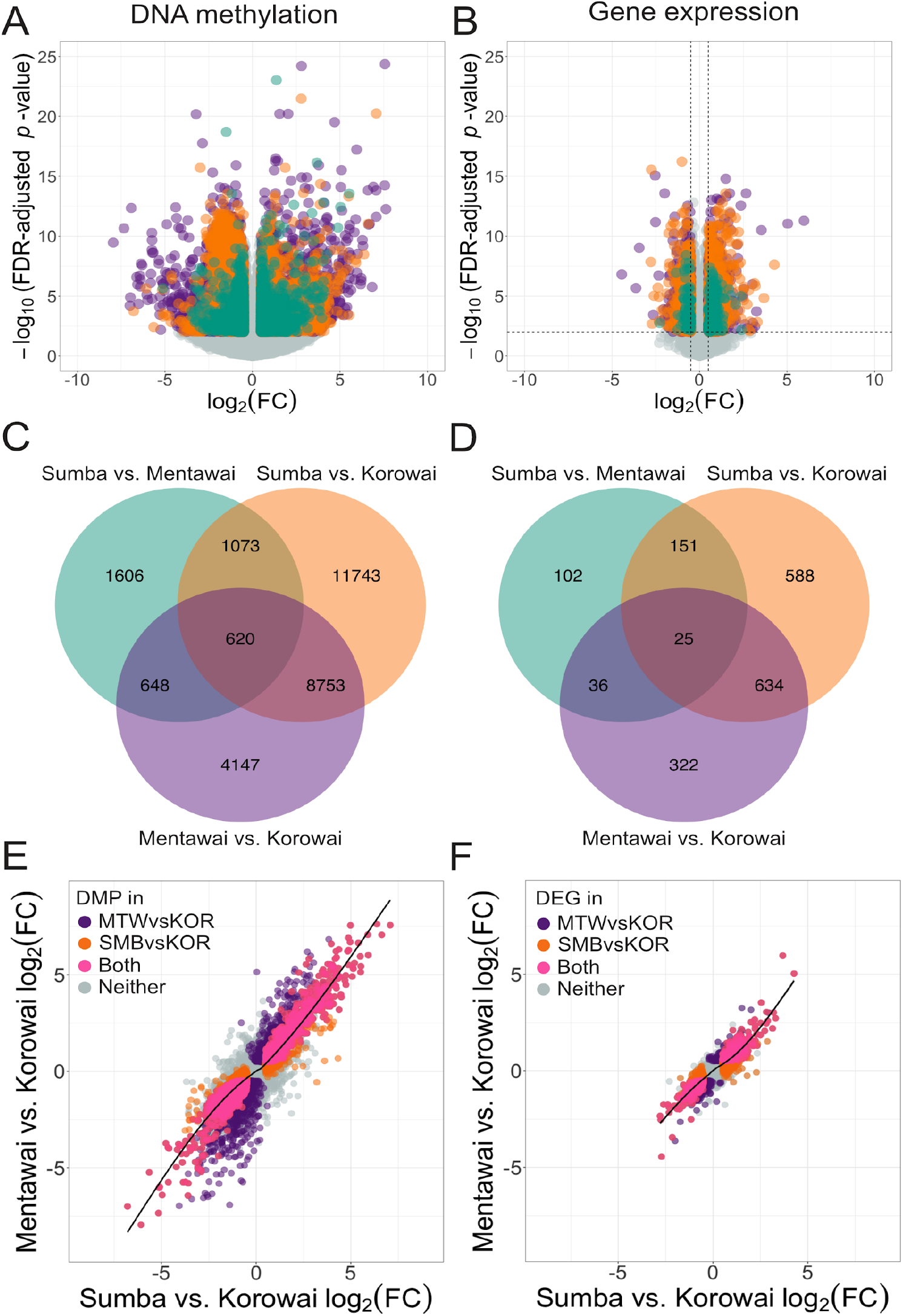
Inter-island differential expression and methylation trends. Volcano plots of (A) differentially methylated probes and (B) differentially expressed genes between Sumba and Mentawai (green), Korowai and Sumba (orange), and Korowai and Mentawai (purple). Venn diagrams of DMPs (C) and DEGs (D) overlapping between different pairwise comparisons at an FDR-adjusted *p* value ≤ 0.01 and an absolute log_2_(FC) ≥ 0.5. Relationship between the log2(FC) of each probe (E) and gene (F) between Mentawai vs. Korowai and Sumba vs. Korowai. Probes and genes that were DMP or DEG between Mentawai and Korowai (purple), Sumba and Korowai (orange), or both comparisons (pink) are indicated. Smoothed conditional means based on generalized additive models are presented with 95% confidence intervals.

There is substantial overlap in signals between either Sumba or Mentawai versus Korowai (Figure 2C, 2D). For instance, 45.35% of DEGs between Sumba and Korowai are also differentially expressed between Mentawai and Korowai; the same is true of 42.24% of DMPs between Sumba and Korowai. DEGs and DMPs between Sumba and Mentawai, however, have poor overlap with the other inter-island comparisons, and are generally limited in number. This suggests that many of the signals we identify are driven by the Korowai data, and by some degree of homogeneity across Sumba and Mentawai. Indeed, comparisons involving Korowai routinely identify an order of magnitude more DEGs and DMPs. Furthermore, we find substantial agreement in both the magnitude and direction of effect between DEGs and DMPs across both comparisons involving Korowai, (Figure 2E, 2F; generalized additive model of the form (y ~ s(x, bs = “cs”)); methylation deviance explained by model = 64.6%,*p* < 2×10^−16^;. expression featuring either Sumba or Mentawai, regardless of whether we focus on methylation or expression differences (Supplementary Figure 5).

### Differentially expressed genes are enriched for immune function and Denisovan introgression

We tested for enrichment of DEGs and DMPs against Gene Ontology (GO^38^) and Kyoto Encyclopedia of Genes and Genomes (KEGG^39^) pathways to detect functional enrichment between island populations. Overlapping enriched GO categories and KEGG pathways (adjusted *p* < 0.05; full tables of results for all comparisons are provided as Supplementary Tables 4-7) in comparisons between both Mentawai or Sumba versus the Korowai include functions related to the adaptive immune response, malaria response, and nervous system function (Supplementary Figure 6). However, DEGs between Mentawai and Sumba were enriched for GO terms related to neurogenesis and the nervous system with no enriched KEGG pathways. Similar testing for enrichment on DMPs shows various categories, which include terms mostly related to neurogenesis, the nervous system, and sensory perception, and which partly overlap with categories enriched in DEGs, although biological interpretation of these terms is not straightforward.

Finally, because the island of New Guinea has the highest levels of Denisovan introgression worldwide (up to 5%^6^), we asked whether any of the genes differentially expressed between the Korowai (high Papuan ancestry) and Mentawai (no Papuan ancestry), or the Korowai and Sumbanese (intermediate Papuan ancestry) fell within high confidence introgressed Denisovan tracts, on the basis of our previous data^6^. A total of 265 DEGs (considering all comparisons) overlap high confidence introgressed Denisovan haplotype blocks in New Guinea^6^. High-frequency introgressed genes in our DEGs includes *FAHD2B* (introgressed at 65% frequency in New Guinea; DE between Sumba and West Papua (*p* = 0.005), and Mentawai and West Papua (*p* = 8.8×10^−7^), and multiple genes related to immunity and antiviral response, such as *CXCR6* (20% frequency in New Guinea^40^) and *GBP1/3/4* (19% frequency in New Guinea^41,42^).

Since calling Denisovan-introgressed genes as differentially expressed depends on both the magnitude of the expression change and the introgressed allele’s frequency, the likelihood cannot be easily predicted *a priori.* Therefore, we examined the distribution of introgressed allele frequencies in New Guinea for all DEGs in our data, and asked whether these differ between our three inter-island comparisons. If Denisovan introgression is contributing to expression differences between the three sampling sites, we expect that genes that are differentially expressed between the Korowai and the other two groups will have generally higher allele frequencies than genes that are DE between the Sumbanese and the Mentawai. Indeed, we observe no difference in allelic frequencies for genes that are DE between both Sumba and West Papua, and Mentawai and West Papua (t-test *p* = 0.946), but observe higher frequencies in DEG between Sumba and West Papua, or Mentawai and West Papua, than between Sumba and Mentawai *(p* = 0.035 and 0.034, respectively), suggesting that Denisovan introgression may impact the expression levels of some genes.

### Methylation changes are associated with changes in gene expression in a subset of genes

To further explore the relationship between DNA methylation and gene expression, we asked how much of the variation we observe in gene expression levels can be attributed to variation in DNA methylation levels. We searched for regions of functional DNA methylation by identifying instances of significant negative correlation between gene expression levels and *cis*-promoter methylation. We identified 1,292 probe clusters associated with 1,261 genes (9.72% of all genes under investigation) where expression level was predicted by nearby CpG methylation (Figure 3A, supplementary table 8). We compared the genes identified in this analysis with the DMPs and DEGs detected in the between-island comparisons, and find that 153 genes (10.94% of DEGs) in the comparison between Korowai and Sumba, 113 genes (11.11%) between Korowai and Mentawai, and 12 genes (3.83%) between Sumba and Mentawai have expression levels associated with significant methylation changes at nearby CpGs; these include genes like *SIGLEC7* (Figure 3B), which is involved in antigen presentation and natural killer (NK) cell–dependent tumor immunosurveillance^43^. *SIGLEC7* and other *SIGLEC* family genes are also potential immunotherapeutic targets against cancer^44^. These results confirm the relationship between DNA methylation and gene expression, and suggest a possible role for differential DNA methylation in shaping the patterns of differential gene expression between these populations. There are five enriched KEGG pathways, all broadly involved in immune interactions (Supplementary Table 9), including natural killer cell-mediated cytotoxicity.

**Figure 3.**
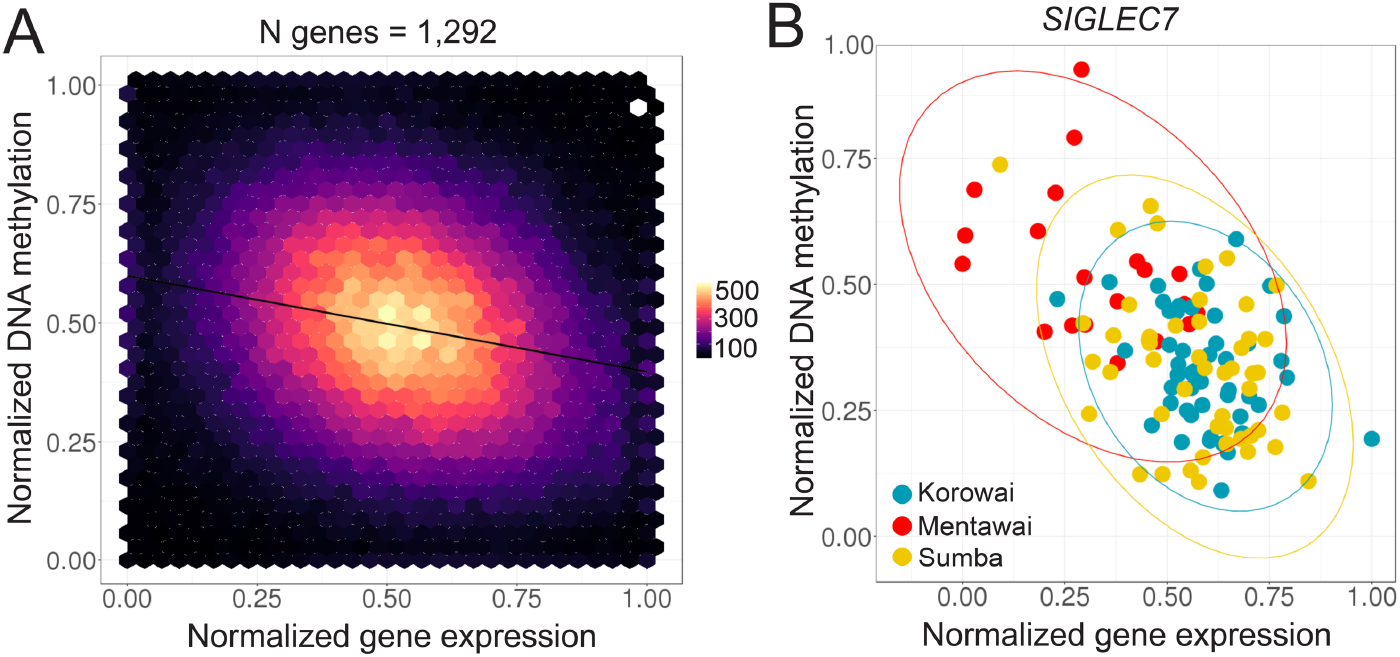
Association between methylation and gene expression levels. (A) Relationship between probe cluster DNA methylation and gene expression levels among the 1,292 probe clusters and associated genes identified by MethylMix. (B) Example of a single gene, *SIGLEC7,* which is both differentially expressed and differentially methylated between Sumbanese and the Korowai.

### Inter-island differences are primarily driven by a subset of villages

While the three island populations differ substantially in terms of genetic composition, we have previously shown that there is a high degree of genetic similarity within islands^13^. Therefore, we may expect that intra-island differences in either DNA methylation or gene expression profiles, if they exist, are likely to reflect local environmental differences^45^. To test this hypothesis, we took advantage of the fact that we collected samples across multiple villages in both Sumba and Mentawai.

PCA captured differences between villages at both the expression and methylation level. For instance, PC1 of the DNA methylation data captures varying degrees of separation at both the intra- and interisland level. Neither the two Sumba villages, Wunga and Anakalang, or the two Mentawai villages, Taileleu and Madobag, are separated by the first PCs, confirming our previous observations of limited differentiation within islands. Between islands, however, PC1 separates the villages of Wunga and Taileleu (Tukey HSD, *p* = 0.001; Supplementary Table 10), Wunga and Madobag (*p* = 0.012), and Anakalang and Taileleu (*p* = 0.017), but not Anakalang and Madobag (*p* = 0.101). Of the two Mentawai villages, Taileleu is clearly separated from Korowai by PC1 (*p* = 1.9×10^−5^), while Madobag is only weakly separated from Korowai (*p* = 0.021); in Sumba, PC1 clearly separates Wunga and Korowai (*p* = 0.003), but separates Anakalang and Korowai only weakly (*p* = 0.033). In the expression data, PC1 separates Mentawai (*p* = 1.0×10^−7^) and Sumba (*p* = 1.5×10^−6^) from Korowai, and PC2 separates Sumba from Mentawai (*p* = 1.6×10^−4^). When examining the villages, PC1 separates the Korowai village from the two Mentawai villages Madobag (*p* = 0.029) and Taileleu (*p* < 1.0×10^−10^) and the Sumba villages Wunga (*p* = 1.0×10^−7^) and Anakalang (*p* = 4.0×10^−4^). PC2 further separates Wunga (*p* = 0.0035) and Anakalang (*p* = 0.039) from Taileleu.

We then repeated our differential expression and methylation analyses between villages. At a log2 FC threshold of 0.5 and an FDR of 1%, we are able to recapitulate the main findings of our island-level analyses, although additional trends emerge (Figure 4, Supplementary Figure 7). Detectable differences between villages in the same island are small, with only 71 DMPs and 51 DEGs between the two Mentawai villages of Madobag and Taileleu, and 21 DMPs and 1 DEG, *IDO1* (a modulator of T-cell behavior and marker of immune activity^46^; *p* = 0.007, log2 FC = −1.48), between the Sumbanese villages of Wunga and Anakalang, echoing their limited separation in the PCA. Similarly, we find low numbers of DEGs and DMPs across all comparisons involving Sumba and Mentawai (Figure 4), again recapitulating the observations we made at the island level (Figure 2). Overall, there appears to be high concordance between genes identified as DE at the island and village level (Supplementary Figure 8), with a high degree of correlation between village- and island-level results, as expected (Supplementary Table 11). However, when comparing villages within islands, we identified substantially more DMPs and DEGs between Taileleu and Korowai (9,631 and 1,157, respectively) than between Madobag and Korowai (7,282 and 486, respectively). Similarly, we identified more DMPs and DEGs between Wunga and Korowai (24,557 and 1,617, respectively) than between Anakalang and Korowai (18,663 and 863, respectively).

**Figure 4.**
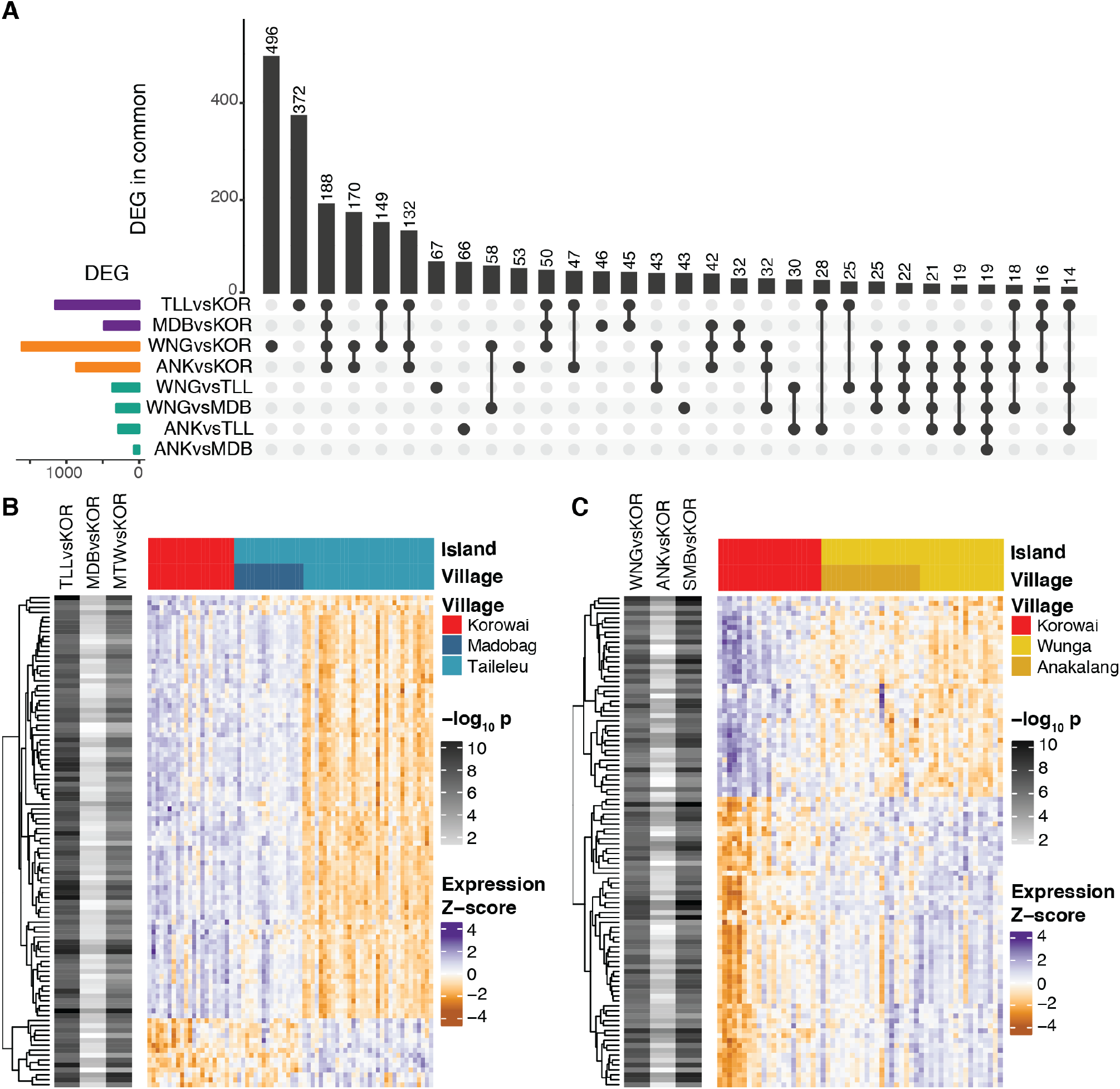
Differential gene expression trends at the village level partially reflect inter-island trends. (A) Sharing of village-level DEG signal across all possible inter-island contrasts. (B) Top 100 DEGs between Taileleu and the Korowai that are not DE between Madobag and the Korowai. (C) Top 100 DEGs between Wunga and the Korowai that are not DE between Anakalang and the Korowai.

We thus focused on genes that exhibit discordant patterns between the villages in an island. DEGs between Taileleu and Korowai, but not between Madobag and Korowai (Figure 4B), tend to have similar expression profiles in Madobag and Korowai, whereas DEGs between Wunga and Korowai but not between Anakalang and Korowai (Figure 4C) seem to be expressed at an intermediate level in Anakalang. These differences are not correlated with known technical confounders such as differences in RNA quality or in variability within villages (Supplementary Figure 9). Indeed, their presence in both the DNA methylation and RNA sequencing results argues against sample processing artifacts. In order to confirm that these patterns were not driven by differences in sample size, we randomly subsampled each village to 10 individuals and repeated DEG testing 10^3^ times. There are consistently more DEGs between Wunga and Korowai than Anakalang and Korowai (t-test *p* < 10^−20^) as well as between Taileleu and Korowai than between Madobag and Korowai (*p* < 10^−20^). In turn, this suggests that they may be driven by interactions between genetics and differences in the local environment at each sampling site, although a comparison of rainfall and mean monthly temperatures across all five sites did not support these factors as drivers (Supplementary Figure 10). On the whole, our results highlight the importance of detailed data collection and thorough sampling from regions spanning diverse genomic and environmental clines, if we are to elucidate gene-by-environment interactions.

## Discussion

Although Island Southeast Asia accounts for nearly 6% of the world’s population, and contains substantial ethnic and genetic diversity^13^, genomic characterisation of this region lags drastically behind other regions of the world. The first regional large-scale set of publicly available human whole genome sequences were published in 2019^6^; to our knowledge there is only one study of gene expression from the region, of patients with malaria from the northern tip of Sulawesi^7^. In contrast, our work represents the first characterization of gene expression and DNA methylation levels across self-reported healthy individuals from geographically and genetically distinct populations in Indonesia, and more broadly from Island Southeast Asia. We have surveyed three sites with genetically distinct populations, spanning the Asian/Papuan genetic cline that characterises human diversity in the region, and we also sampled multiple villages in two of the islands (Sumba and Mentawai). Our study design purposefully allows us to explore both genetic (primarily between islands) and environmental (both between and within island) contributions to expression and methylation differences, a result that is further highlighted in our intervillage analysis, where we observe some small-scale village-specific effects (Figure 4).

Indeed, while we find differentially expressed genes and differentially methylated CpGs in most location comparisons (Figure 2), the most numerous, reproducible and largest effect changes were found when comparing either the Sumbanese or Mentawai with the Korowai. Many of these results feature genes involved in immune function, suggesting a potentially adaptive response to local environmental pressures. For example, beyond consistent enrichment for immune-associated GO and KEGG terms, the top 20 strongest DEG signals between the Mentawai and the Korowai include genes involved in antigen presentation in both innate and adaptive immune cells (*MARCO* and *SIGLEC7,* respectively; *MARCO p* = 2.7×10^−14^; *SIGLEC7 p* = 9.7×10^−14^; these genes are also differentially expressed between Sumbanese and the Korowai (*MARCO p* = 4.2×10^−10^; *SIGLEC7 p* = 4.9×10^−12^; supplementary figure 11). Polymorphisms within MARCO, which is expressed on the surface of macrophages, have been repeatedly shown to associate with susceptibility of infection by *Mycobacterium tuberculosis* and *Streptococcus pneumoniae* in multiple populations worldwide^47–50^; some of these variants have been subsequently shown to have a direct impact on antigen binding^51^. Our MethylMix analyses identify differences in *SIGLEC7* expression as being driven, at least in part, by methylation differences in its promoter region (Figure 4C).

In the absence of whole genome data from our samples, it is challenging to identify whether these signals are also associated with selective signals at the DNA level or driven entirely by environmental differences; neither of these genes has been identified in previous scans of Denisovan introgressions. However, both we and others have previously shown that introgressed Denisovan tracts on the island of New Guinea are enriched for immune genes^6,52^, similar to the contributions of Neandertals to non-African genomes^53,54^. Indeed, our data suggest that Denisovan introgression in New Guinea may be impacting gene expression levels in the Korowai. More broadly, immune challenges have exerted some of the strongest selective forces on humans throughout our species’ history^11^; transmissible diseases endemic in Indonesia range from malaria (both *P. falciparum* and *P. vivax*)^8^ to infections by multiple helminth species and other understudied tropical diseases^2^. Tuberculosis remains a major health concern in the region, with the World Health Organisation reporting nearly half a million new cases in 2017^55^.

Others have sought to characterise the interplay between genetic and environmental contributions to either expression or methylation levels across limited geographic scales. A study of approximately 1,000 individuals drawn from a founder population in Quebec demonstrated that gene-by-environment interactions – specifically, with air pollution levels – drastically impacted measurements of gene expression in blood, overpowering the effects of genetic relatedness^45^. Equivalent high-resolution Indonesian data are unavailable, and our attempts to associate differences in expression or methylation across small geographic scales by using WorldClim data were inconclusive. Unfortunately, it remains difficult to characterize granular levels of regional heterogeneity in disease burden and infection type, yet our results suggest pressures shaping immune response in Indonesia vary at the local level.

A different study of DNA methylation across rainforest hunter-gatherer and farmer populations in Central Africa showed that methylation captures both population history and current lifestyle practices. However, these two factors impact non-overlapping sets of genes, with differences at immune genes associated with a group’s present-day habitat as well as genomic signals of past positive selection^45^. We observe similar trends here; the Korowai occupy an ecological niche akin to that of African rainforest huntergatherers, whereas the inhabitants of Sumba and Mentawai are village-based agriculturalists. Sumba in particular is host to a network of traditional communities derived largely from pre-existing Papuans, who first arrived on the island ~50,000 years ago, and incoming Asian farming cultures, that reached the island ~4,000 years ago^14^. Today, Sumba retains a low population density and little contact between villages, as reflected in its extensive linguistic diversity^56^. This has resulted in small, isolated populations of a few hundred to a few thousand individuals that can be identified genetically between villages roughly 10 km apart^14^, making it a near unique study system for examining gene by environment interactions.

As we move further into the age of personalised and genomic medicine, understanding how genetics and other molecular phenotypes drive disease risk across diverse populations is of crucial importance to ensure benefits are equitably distributed. Already there has been a dramatic expansion of genomic-based tests that are being deployed to identify the risk of disease. However, these tests are largely built using European cohorts and have proven difficult to translate to non-European populations^57–59^. Even within homogeneous populations, environmental factors can have marked effects on gene expression measurements, and on the interpretability of genomic-based tests of disease risk^60^, highlighting a secondary risk of such biased European sampling: limiting not only the genomic diversity under study, but the environmental diversity as well, to general detriment. This study provides a valuable first step in the characterization of the processes shaping gene expression changes in Island Southeast Asia.

## Acknowledgements

We especially thank all of our study participants and the Eijkman Institute field survey team, without whom this work would not have been possible. We thank Nicolas Brucato (Université de Toulouse Midi-Pyrénées), Christine Wells (University of Melbourne), Davide Vespasiani (University of Melbourne) and Isabella Apriyana (Australian National University) for valuable discussion. This study was supported by a National Science Foundation Grant SES 0725470 and a Singapore Ministry of Education Tier II Grant MOE2015-T2-1-127 to JSL, an NTU Presidential Postdoctoral Fellowship to GSJ, an NTU Complexity Institute Individual Fellowship to PK, and a Royal Society of New Zealand Marsden Grant 17-MAU-040 to MPC and IGR. HN was supported by an ASU Center for Evolution and Medicine postdoctoral fellowship and the Marcia and Frank Carlucci Charitable Foundation postdoctoral award from the Prevent Cancer Foundation. KSB was supported by a Melbourne Graduate Research Scholarship. MPC was supported by a University of Melbourne Miegunyah Distinguished Visiting Professor fellowship.

## Supplementary Materials

Supplementary materials include 11 tables and 11 figures:

Supplementary table 1: Sample metadata

Supplementary table 2: Sample sequencing information

Supplementary table 3: Summary of DEG/DMP/DMR testing at various thresholds

Supplementary table 4: GO enrichment testing results for DEGs

Supplementary table 5: KEGG enrichment testing results for DEGs

Supplementary table 6: GO enrichment testing results for DMPs

Supplementary table 7: KEGG enrichment testing results for DMPs

Supplementary table 8: List of significant MethylMix clusters

Supplementary table 9: KEGG enrichment testing for MethylMix-associated genes

Supplementary table 10: ANOVA on PCA and covariates

Supplementary table 11: Spearman correlation between village and island level across both DEG and DMP tests

Supplementary figure 1: Clustering of the gene expression data before and after batch correction

Supplementary figure 2: Distribution of Spearman’s pairwise correlation (rho) values across all levels of the RNA-sequencing data

Supplementary figure 3: Clustering of the DNA methylation data before and after batch correction

Supplementary figure 4: Distribution of Spearman’s pairwise correlation (rho) values across all levels of the DNA methylation data.

Supplementary figure 5: Relationship between the log2(FC) of probes and genes across island-level comparisons.

Supplementary figure 6: Shared GO terms between Sumbanese and the Korowai and the Mentawai and the Korowai

Supplementary figure 7: Sharing of village-level DMP signal across all possible inter-island contrasts.

Supplementary figure 8: Sharing of DE signals at the island and village levels

Supplementary figure 9: Distribution of coefficients of variation (CoV) across villages

Supplementary figure 10: Monthly climate fluctuations across the five main village sampling sites.

Supplementary figure 11: log2 CPM values across all samples for (A) *MARCO* and (B) *SIGLEC7.*

## References

1. Popejoy, A.B., and Fullerton, S.M. (2016). Genomics is failing on diversity. Nature 538, 161–164.

2. Horton, R. (2016). Offline: Indonesia—unravelling the mystery of a nation. Lancet 387, 830.

3. 1000 Genomes Project Consortium, Auton, A., Brooks, L.D., Durbin, R.M., Garrison, E.P., Kang, H.M., Korbel, J.O., Marchini, J.L., McCarthy, S., McVean, G.A., et al. (2015). A global reference for human genetic variation. Nature 526, 68–74.

4. Mallick, S., Li, H., Lipson, M., Mathieson, I., Gymrek, M., Racimo, F., Zhao, M., Chennagiri, N., Nordenfelt, S., Tandon, A., et al. (2016). The Simons Genome Diversity Project: 300 genomes from 142 diverse populations. Nature 538, 201–206.

5. Pagani, L., Lawson, D.J., Jagoda, E., Mörseburg, A., Eriksson, A., Mitt, M., Clemente, F., Hudjashov, G., DeGiorgio, M., Saag, L., et al. (2016). Genomic analyses inform on migration events during the peopling of Eurasia. Nature 538, 238–242.

6. Jacobs, G.S., Hudjashov, G., Saag, L., Kusuma, P., Darusallam, C.C., Lawson, D.J., Mondal, M., Pagani, L., Ricaut, F.-X., Stoneking, M., et al. (2019). Multiple Deeply Divergent Denisovan Ancestries in Papuans. Cell 177, 1010–1021.e32.

7. Yamagishi, J., Natori, A., Tolba, M.E.M., Mongan, A.E., Sugimoto, C., Katayama, T., Kawashima, S., Makalowski, W., Maeda, R., Eshita, Y., et al. (2014). Interactive transcriptome analysis of malaria patients and infecting Plasmodium falciparum. Genome Res. 24, 1433–1444.

8. Elyazar, I.R.F., Hay, S.I., and Baird, J.K. (2011). Malaria distribution, prevalence, drug resistance and control in Indonesia. Adv. Parasitol. 74, 41–175.

9. R. Tedjo Sasmono, Rama Dhenni, Benediktus Yohan, Paul Pronyk, Sri Rezeki Hadinegoro, Elizabeth Jane Soepardi, Chairin Nisa Ma’roef, Hindra I. Satari, Heather Menzies, William A. Hawley, et al. (2018). Zika Virus Seropositivity in 1–4-Year-Old Children, Indonesia, 2014. Emerging Infectious Disease Journal 24, 1740.

10. Suryanto, Plummer, V., and Boyle, M. (2017). Healthcare System in Indonesia. Hosp. Top. 95, 82–89.

11. Quintana-Murci, L. (2019). Human Immunology through the Lens of Evolutionary Genetics. Cell 177, 184–199.

12. Cox, M.P., Karafet, T.M., Lansing, J.S., Sudoyo, H., and Hammer, M.F. (2010). Autosomal and X-linked single nucleotide polymorphisms reveal a steep Asian-Melanesian ancestry cline in eastern Indonesia and a sex bias in admixture rates. Proc. Biol. Sci. 277, 1589–1596.

13. Hudjashov, G., Karafet, T.M., Lawson, D.J., Downey, S., Savina, O., Sudoyo, H., Lansing, J.S., Hammer, M.F., and Cox, M.P. (2017). Complex Patterns of Admixture across the Indonesian Archipelago. Mol. Biol. Evol. 34, 2439–2452.

14. Cox, M.P., Hudjashov, G., Sim, A., Savina, O., Karafet, T.M., Sudoyo, H., and Lansing, J.S. (2016). Small Traditional Human Communities Sustain Genomic Diversity over Microgeographic Scales despite Linguistic Isolation. Mol. Biol. Evol. 33, 2273–2284.

15. Babraham Bioinformatics – FastQC A Quality Control tool for High Throughput Sequence Data. http://www.bioinformatics.babraham.ac.uk/projects/fastqc/.

16. Bolger, A.M., Lohse, M., and Usadel, B. (2014). Trimmomatic: a flexible trimmer for Illumina sequence data. Bioinformatics 30, 2114–2120.

17. Dobin, A., Davis, C.A., Schlesinger, F., Drenkow, J., Zaleski, C., Jha, S., Batut, P., Chaisson, M., and Gingeras, T.R. (2013). STAR: ultrafast universal RNA-seq aligner. Bioinformatics 29, 15–21.

18. Liao, Y., Smyth, G.K., and Shi, W. (2014). featureCounts: an efficient general purpose program for assigning sequence reads to genomic features. Bioinformatics 30, 923–930.

19. R Core Team (2017). R: A language and environment for statistical computing (R Foundation for Statistical Computing, Vienna, Austria).

20. Robinson, M.D., and Oshlack, A. (2010). A scaling normalization method for differential expression analysis of RNA-seq data. Genome Biol. 11, R25.

21. Law, C.W., Chen, Y., Shi, W., and Smyth, G.K. (2014). voom: precision weights unlock linear model analysis tools for RNA-seq read counts. Genome Biol. 15, R29.

22. Ritchie, M.E., Phipson, B., Wu, D., Hu, Y., Law, C.W., Shi, W., and Smyth, G.K. (2015). limma powers differential expression analyses for RNA-sequencing and microarray studies. Nucleic Acids Res. 43, e47.

23. Aguirre-Gamboa, R., de Klein, N., di Tommaso, J., Claringbould, A., Vosa, U., Zorro, M., Chu, X., Bakker, O.O., Borek, Z., Ricano-Ponce, I., et al. (2019). Deconvolution of bulk blood eQTL effects into immune cell subpopulations.

24. Durinck, S., Spellman, P.T., Birney, E., and Huber, W. (2009). Mapping identifiers for the integration of genomic datasets with the R/Bioconductor package biomaRt. Nat. Protoc. 4, 1184–1191.

25. Yu, G., Wang, L.-G., Han, Y., and He, Q.-Y. (2012). clusterProfiler: an R package for comparing biological themes among gene clusters. OMICS 16, 284–287.

26. Carlson, M. Genome wide annotation for Human, primarily based on mapping using Entrez Gene identifiers. https://doi.org/10.18129/B9.bioc.org.Hs.eg.db.

27. Froehlich, H. GOSim. https://doi.org/10.18129/B9.bioc.GOSim.

28. Fick, S.E., and Hijmans, R.J. (2017). WorldClim 2: new 1-km spatial resolution climate surfaces for global land areas. International Journal of Climatology 37, 4302–4315.

29. Aryee, M.J., Jaffe, A.E., Corrada-Bravo, H., Ladd-Acosta, C., Feinberg, A.P., Hansen, K.D., and Irizarry, R.A. (2014). Minfi: a flexible and comprehensive Bioconductor package for the analysis of Infinium DNA methylation microarrays. Bioinformatics 30, 1363–1369.

30. Maksimovic, J., Gordon, L., and Oshlack, A. (2012). SWAN: Subset-quantile within array normalization for illumina infinium HumanMethylation450 BeadChips. Genome Biol. 13, R44.

31. Du, P., Kibbe, W.A., and Lin, S.M. (2008). lumi: a pipeline for processing Illumina microarray. Bioinformatics 24, 1547–1548.

32. Phipson, B., Maksimovic, J., and Oshlack, A. (2016). missMethyl: an R package for analyzing data from Illumina’s HumanMethylation450 platform. Bioinformatics 32, 286–288.

33. Geeleher, P., Hartnett, L., Egan, L.J., Golden, A., Raja Ali, R.A., and Seoighe, C. (2013). Gene-set analysis is severely biased when applied to genome-wide methylation data. Bioinformatics 29, 1851–1857.

34. Peters, T.J., Buckley, M.J., Statham, A.L., Pidsley, R., Samaras, K., V Lord, R., Clark, S.J., and Molloy, P.L. (2015). De novo identification of differentially methylated regions in the human genome. Epigenetics Chromatin 8, 6.

35. Gevaert, O. (2015). MethylMix: an R package for identifying DNA methylation-driven genes. Bioinformatics 31, 1839–1841.

36. Cedoz, P.-L., Prunello, M., Brennan, K., and Gevaert, O. (2018). MethylMix 2.0: an R package for identifying DNA methylation genes. Bioinformatics 34, 3044–3046.

37. Reich, D., Patterson, N., Kircher, M., Delfin, F., Nandineni, M.R., Pugach, I., Ko, A.M.-S., Ko, Y.-C., Jinam, T.A., Phipps, M.E., et al. (2011). Denisova admixture and the first modern human dispersals into Southeast Asia and Oceania. Am. J. Hum. Genet. 89, 516–528.

38. The Gene Ontology Consortium (2019). The Gene Ontology Resource: 20 years and still GOing strong. Nucleic Acids Res. 47, D330–D338.

39. Kanehisa, M., and Goto, S. (2000). KEGG: kyoto encyclopedia of genes and genomes. Nucleic Acids Res. 28, 27–30.

40. Paust, S., Gill, H.S., Wang, B.-Z., Flynn, M.P., Moseman, E.A., Senman, B., Szczepanik, M., Telenti, A., Askenase, P.W., Compans, R.W., et al. (2010). Critical role for the chemokine receptor CXCR6 in NK cell-mediated antigen-specific memory of haptens and viruses. Nat. Immunol. 11, 1127–1135.

41. Shenoy, A.R., Kim, B.-H., Choi, H.-P., Matsuzawa, T., Tiwari, S., and MacMicking, J.D. (2007). Emerging themes in IFN-gamma-induced macrophage immunity by the p47 and p65 GTPase families. Immunobiology 212, 771–784.

42. Pilla-Moffett, D., Barber, M.F., Taylor, G.A., and Coers, J. (2016). Interferon-Inducible GTPases in Host Resistance, Inflammation and Disease. J. Mol. Biol. 428, 3495–3513.

43. Jandus, C., Boligan, K.F., Chijioke, O., Liu, H., Dahlhaus, M., Démoulins, T., Schneider, C., Wehrli, M., Hunger, R.E., Baerlocher, G.M., et al. (2014). Interactions between Siglec-7/9 receptors and ligands influence NK cell–dependent tumor immunosurveillance. Journal of Clinical Investigation 124, 1810–1820.

44. Daly, J., Carlsten, M., and O’Dwyer, M. (2019). Sugar Free: Novel Immunotherapeutic Approaches Targeting Siglecs and Sialic Acids to Enhance Natural Killer Cell Cytotoxicity Against Cancer. Frontiers in Immunology 10,.

45. Favé, M.-J., Lamaze, F.C., Soave, D., Hodgkinson, A., Gauvin, H., Bruat, V., Grenier, J.-C., Gbeha, E., Skead, K., Smargiassi, A., et al. (2018). Gene-by-environment interactions in urban populations modulate risk phenotypes. Nat. Commun. 9, 827.

46. Zhai, L., Ladomersky, E., Lenzen, A., Nguyen, B., Patel, R., Lauing, K.L., Wu, M., and Wainwright, D.A. (2018). IDO1 in cancer: a Gemini of immune checkpoints. Cell. Mol. Immunol. 15, 447.

47. Bowdish, D.M.E., Sakamoto, K., Lack, N.A., Hill, P.C., Sirugo, G., Newport, M.J., Gordon, S., Hill, A.V.S., and Vannberg, F.O. (2013). Genetic variants of MARCO are associated with susceptibility to pulmonary tuberculosis in a Gambian population. BMC Medical Genetics 14,.

48. Ma, M.-J., Wang, H.-B., Li, H., Yang, J.-H., Yan, Y., Xie, L.-P., Qi, Y.-C., Li, J.-L., Chen, M.-J., Liu, W., et al. (2011). Genetic variants in MARCO are associated with the susceptibility to pulmonary tuberculosis in Chinese Han population. PLoS One 6, e24069.

49. Dorrington, M.G., Roche, A.M., Chauvin, S.E., Tu, Z., Mossman, K.L., Weiser, J.N., and Bowdish, D.M.E. (2013). MARCO Is Required for TLR2- and Nod2-Mediated Responses to Streptococcus pneumoniae and Clearance of Pneumococcal Colonization in the Murine Nasopharynx. The Journal of Immunology 190, 250–258.

50. Thuong, N.T.T., Tram, T.T.B., Dinh, T.D., Thai, P.V.K., Heemskerk, D., Bang, N.D., Chau, T.T.H., Russell, D.G., Thwaites, G.E., Hawn, T.R., et al. (2016). MARCO variants are associated with phagocytosis, pulmonary tuberculosis susceptibility and Beijing lineage. Genes Immun. 17, 419–425.

51. Novakowski, K.E., Yap, N.V.L., Yin, C., Sakamoto, K., Heit, B., Golding, G.B., and Bowdish, D.M.E. (2018). Human-Specific Mutations and Positively Selected Sites in MARCO Confer Functional Changes. Mol. Biol. Evol. 35, 440–450.

52. Gittelman, R.M., Schraiber, J.G., Vernot, B., Mikacenic, C., Wurfel, M.M., and Akey, J.M. (2016). Archaic Hominin Admixture Facilitated Adaptation to Out-of-Africa Environments. Curr. Biol. 26, 3375–3382.

53. Abi-Rached, L., Jobin, M.J., Kulkarni, S., McWhinnie, A., Dalva, K., Gragert, L., Babrzadeh, F., Gharizadeh, B., Luo, M., Plummer, F.A., et al. (2011). The shaping of modern human immune systems by multiregional admixture with archaic humans. Science 334, 89–94.

54. Dannemann, M., Andrés, A.M., and Kelso, J. (2016). Introgression of Neandertal- and Denisovan-like Haplotypes Contributes to Adaptive Variation in Human Toll-like Receptors. Am. J. Hum. Genet. 98, 22–33.

55. WHO (2019). Tuberculosis country profiles. https://www.who.int/tb/country/data/profiles/en/ (World Health Organization).

56. Lansing, J.S., Cox, M.P., Downey, S.S., Gabler, B.M., Hallmark, B., Karafet, T.M., Norquest, P., Schoenfelder, J.W., Sudoyo, H., Watkins, J.C., et al. (2007). Coevolution of languages and genes on the island of Sumba, eastern Indonesia. Proc. Natl. Acad. Sci. U. S. A. 104, 16022–16026.

57. Martin, A.R., Kanai, M., Kamatani, Y., Okada, Y., Neale, B.M., and Daly, M.J. (2019). Clinical use of current polygenic risk scores may exacerbate health disparities. Nat. Genet. 51, 584–591.

58. Daar, A.S., and Singer, P.A. (2005). Pharmacogenetics and geographical ancestry: implications for drug development and global health. Nat. Rev. Genet. 6, 241–246.

59. Martin, A.R., Gignoux, C.R., Walters, R.K., Wojcik, G.L., Neale, B.M., Gravel, S., Daly, M.J., Bustamante, C.D., and Kenny, E.E. (2017). Human Demographic History Impacts Genetic Risk Prediction across Diverse Populations. Am. J. Hum. Genet. 100, 635–649.

60. Mostafavi, H., Harpak, A., Conley, D., Pritchard, J.K., and Przeworski, M. Variable prediction accuracy of polygenic scores within an ancestry group.

