## Supplementary Figures 1-11 for "Genome-wide DNA methylation and gene expression patterns reflect genetic ancestry and environmental differences across the Indonesian archipelago"

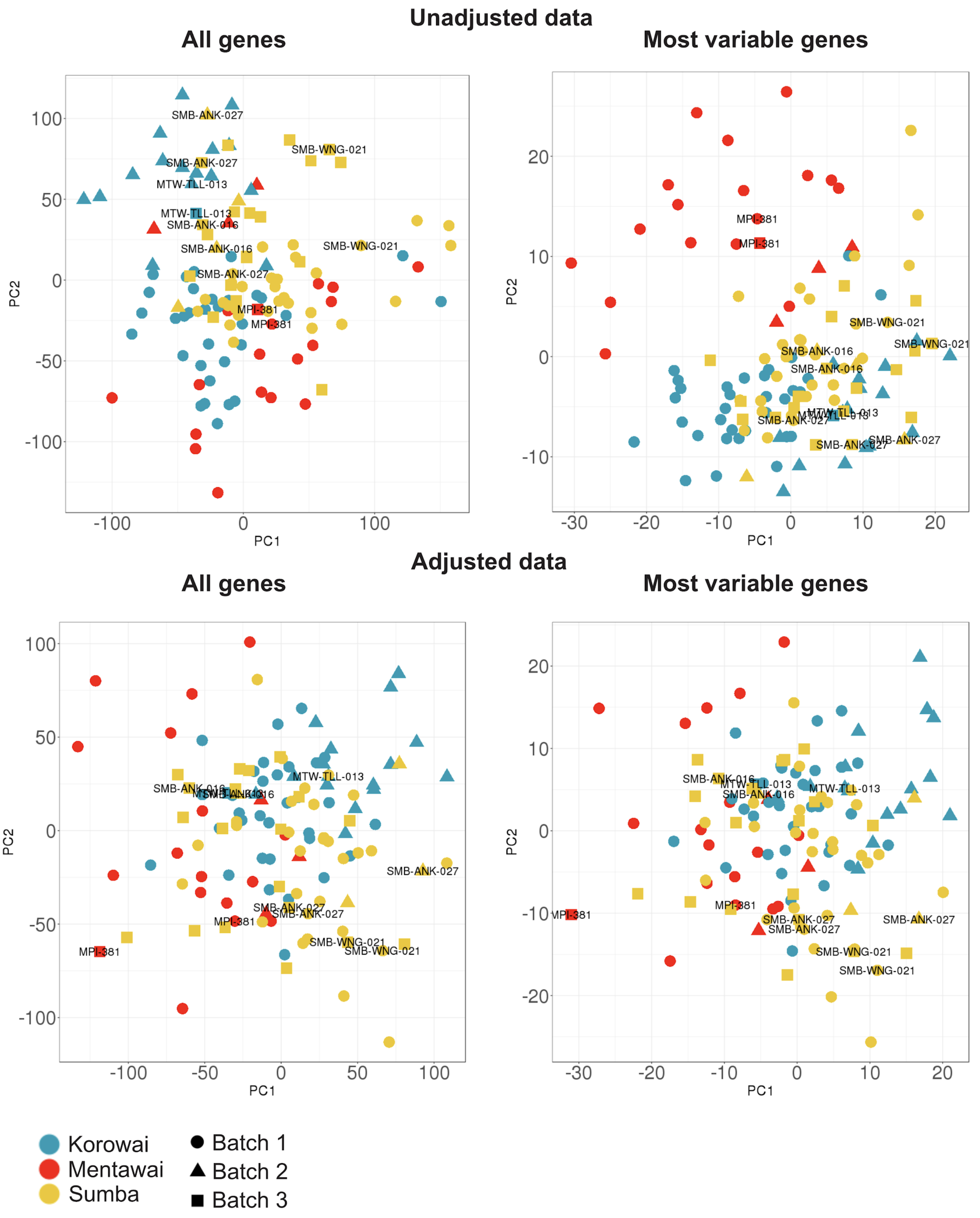


**Supplementary Figure 1** Clustering of the gene expression data before and after batch correction. PCA using most variable genes is based on 1,000 genes exhibiting the largest coefficient of variance between samples. Technical replicates are labeled.

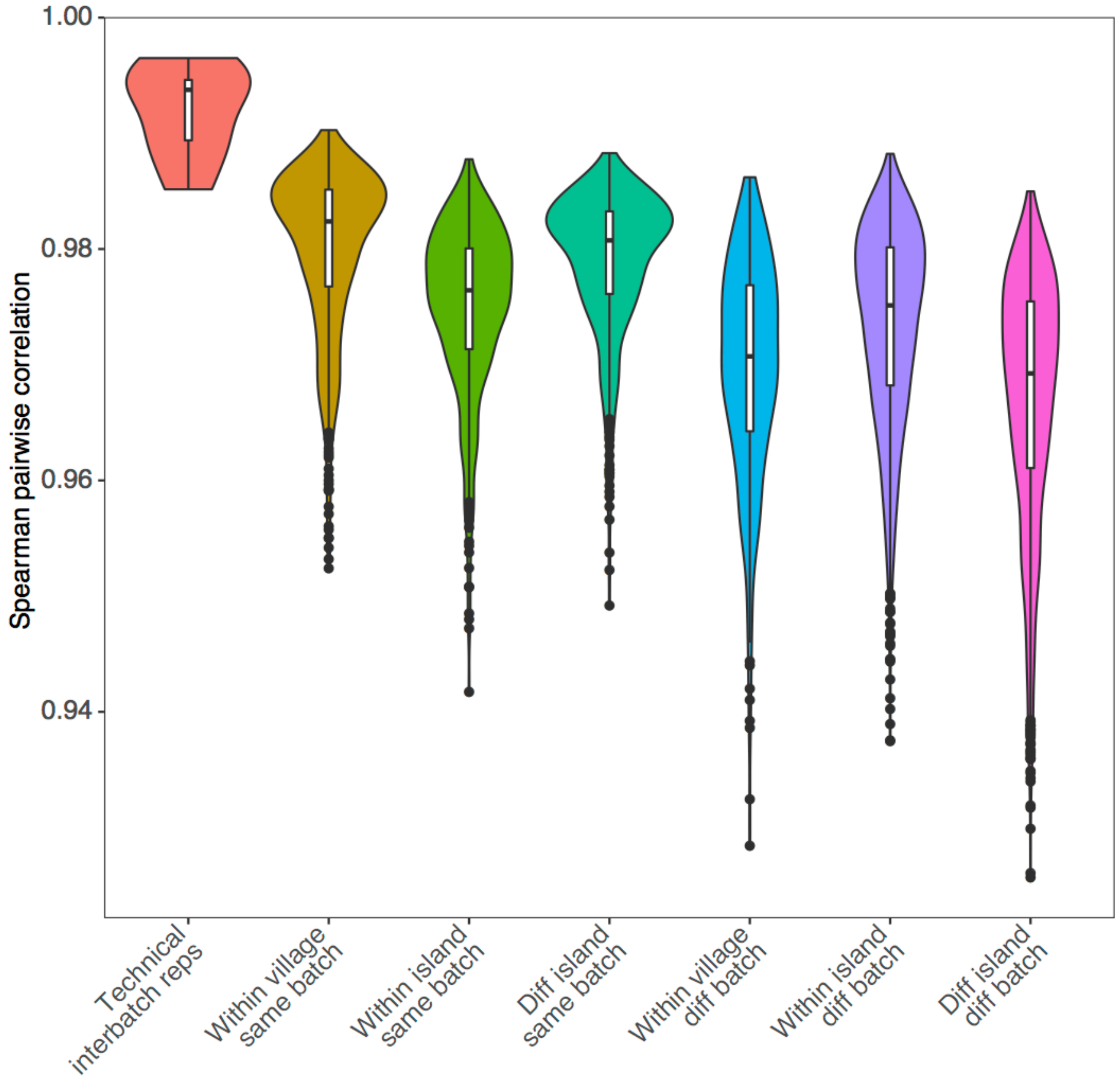


**Supplementary figure 2:** Distribution of Spearman's pairwise correlation (rho) values across all levels of the RNA-sequencing data.

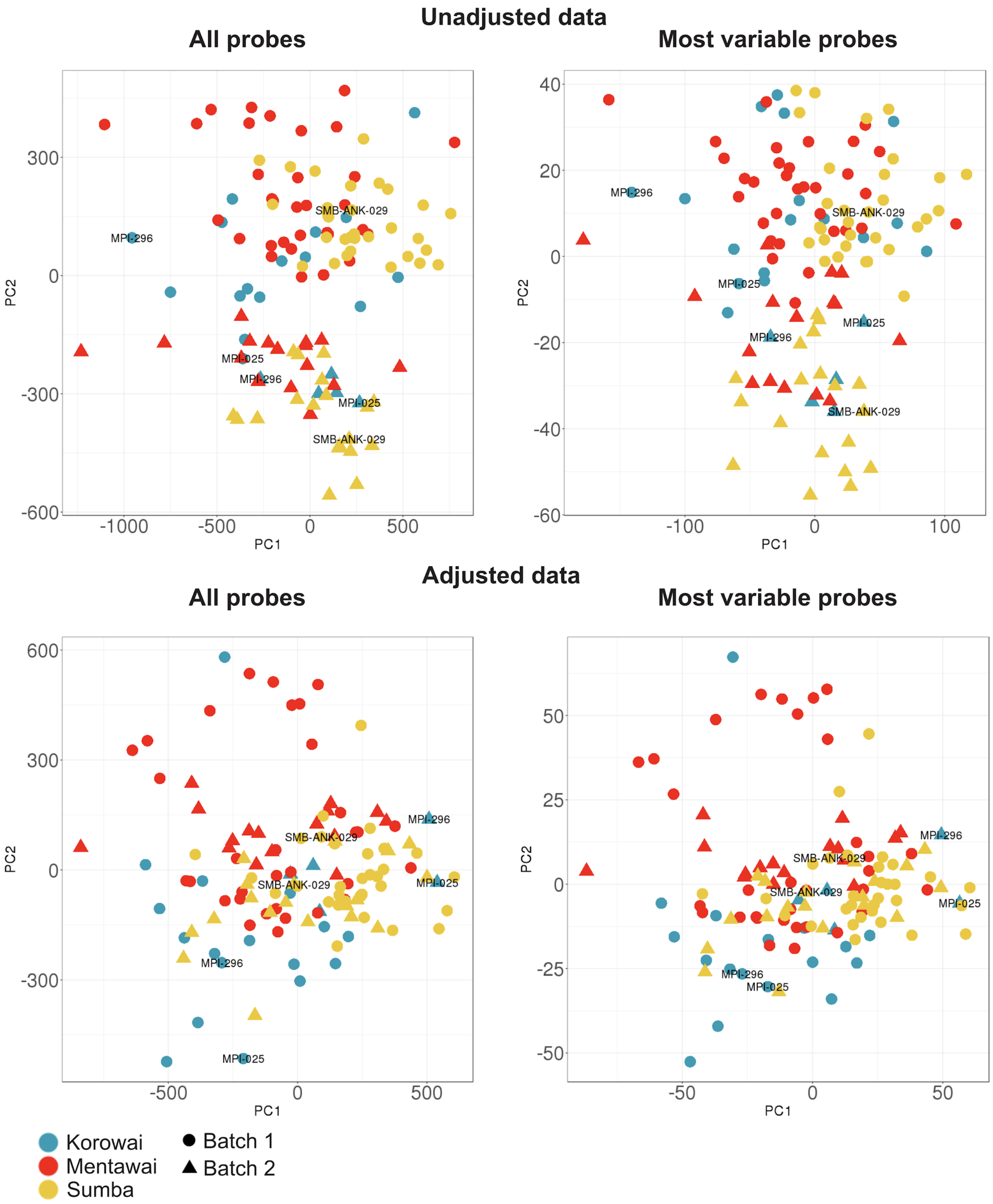


**Supplementary Figure 3:** Clustering of the DNA methylation data before and after batch correction. PCA using most variable probes is based on 10,000 probes exhibiting the largest coefficient of variance between samples. Technical replicates are labeled.

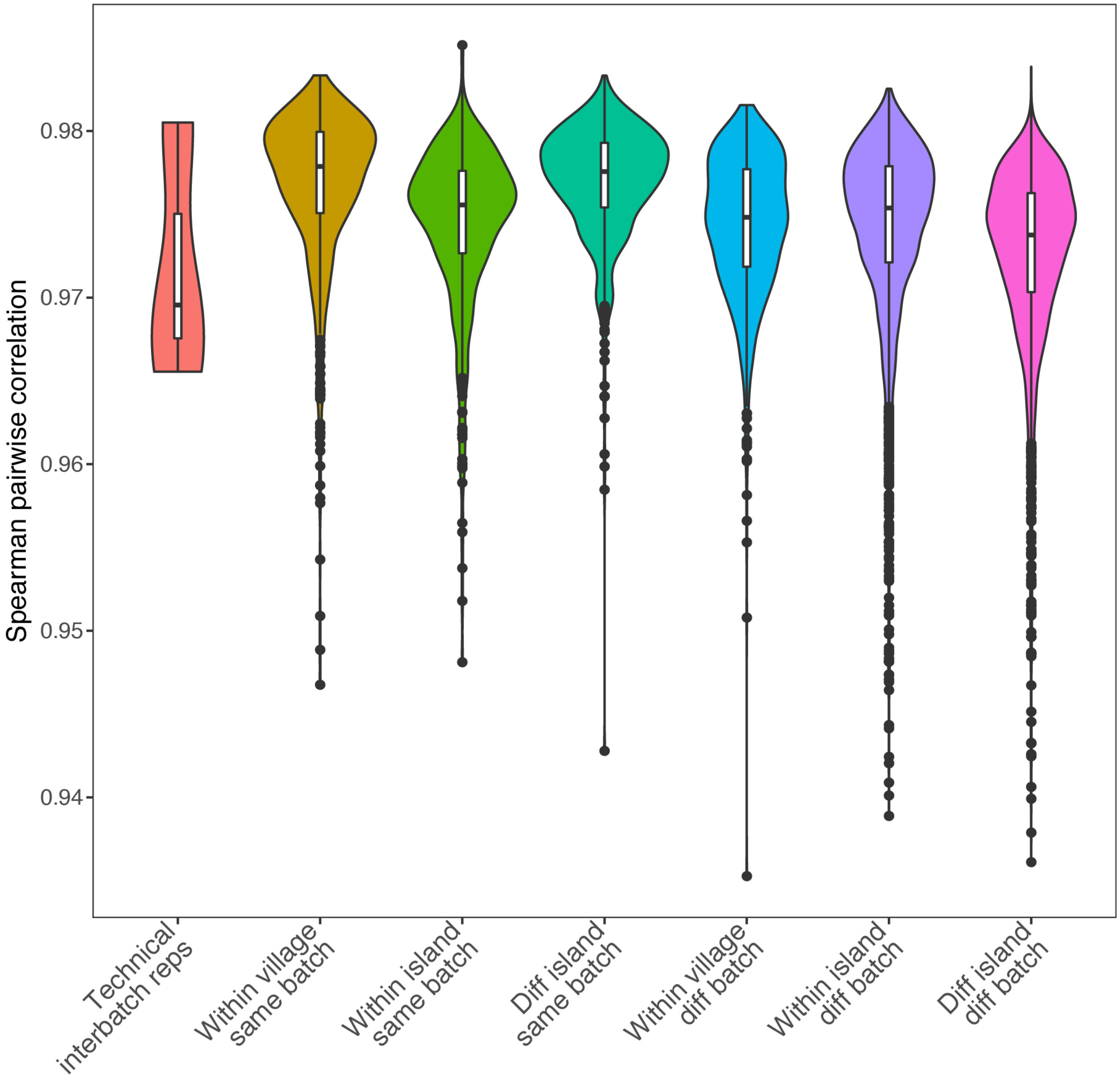


**Supplementary figure 4:** Distribution of Spearman's pairwise correlation (rho) values across all levels of the DNA methylation data.

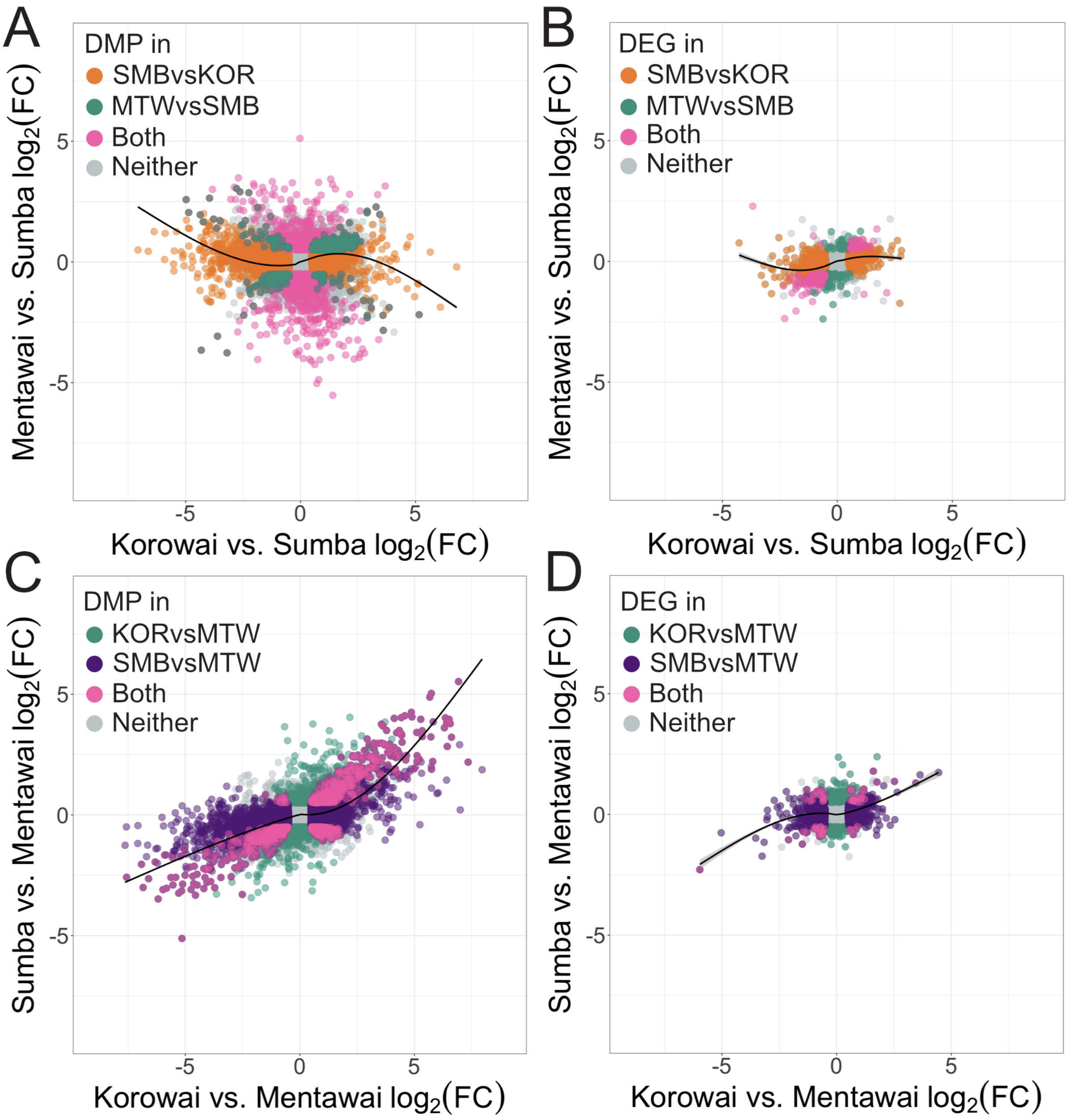


**Supplementary Figure 5:** Relationship of $\log_2(FC)$ in the probe methylation levels (A,C) and gene expression levels (B,D) between the Korowai vs. Sumba and Mentawai vs. Sumba comparisons (A,B) and between the Korowai vs. Mentawai and Sumba vs. Mentawai comparisons (C,D).

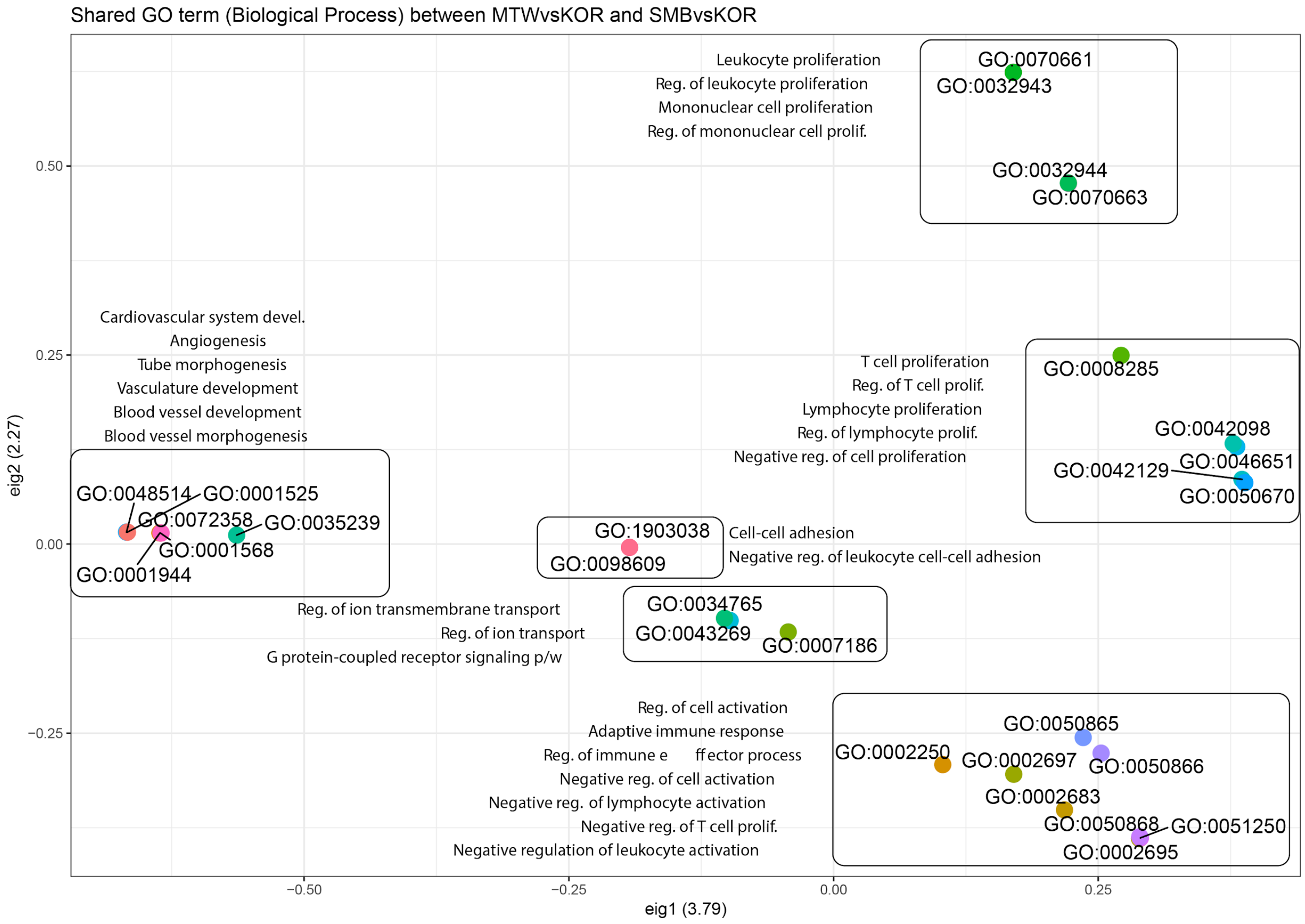


**Supplementary figure 6:** Shared GO terms between Sumbanese and the Korowai and the Mentawai and the Korowai. Term similarity was established by using GOSim and captures a high degree of term sharing and similarities in immunity-associated terms across the two contrasts.

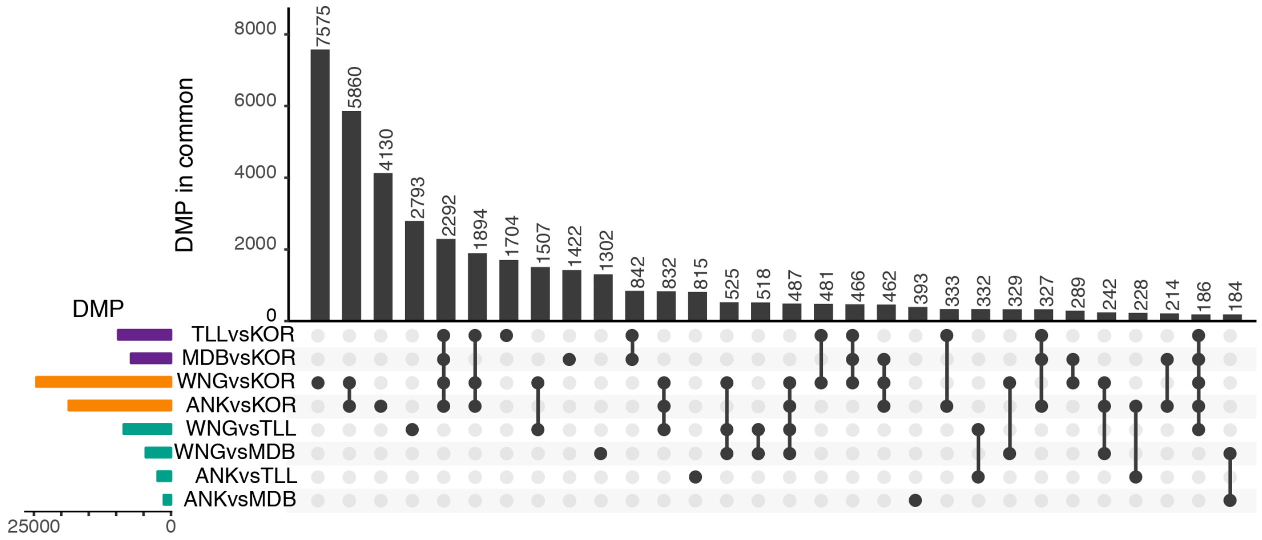


**Supplementary Figure 7:** Sharing of village-level DMP signal across all possible inter-island contrasts. All results are thresholded at an FDR of 1% and an absolute $\log_2 FC$ greater than 0.5

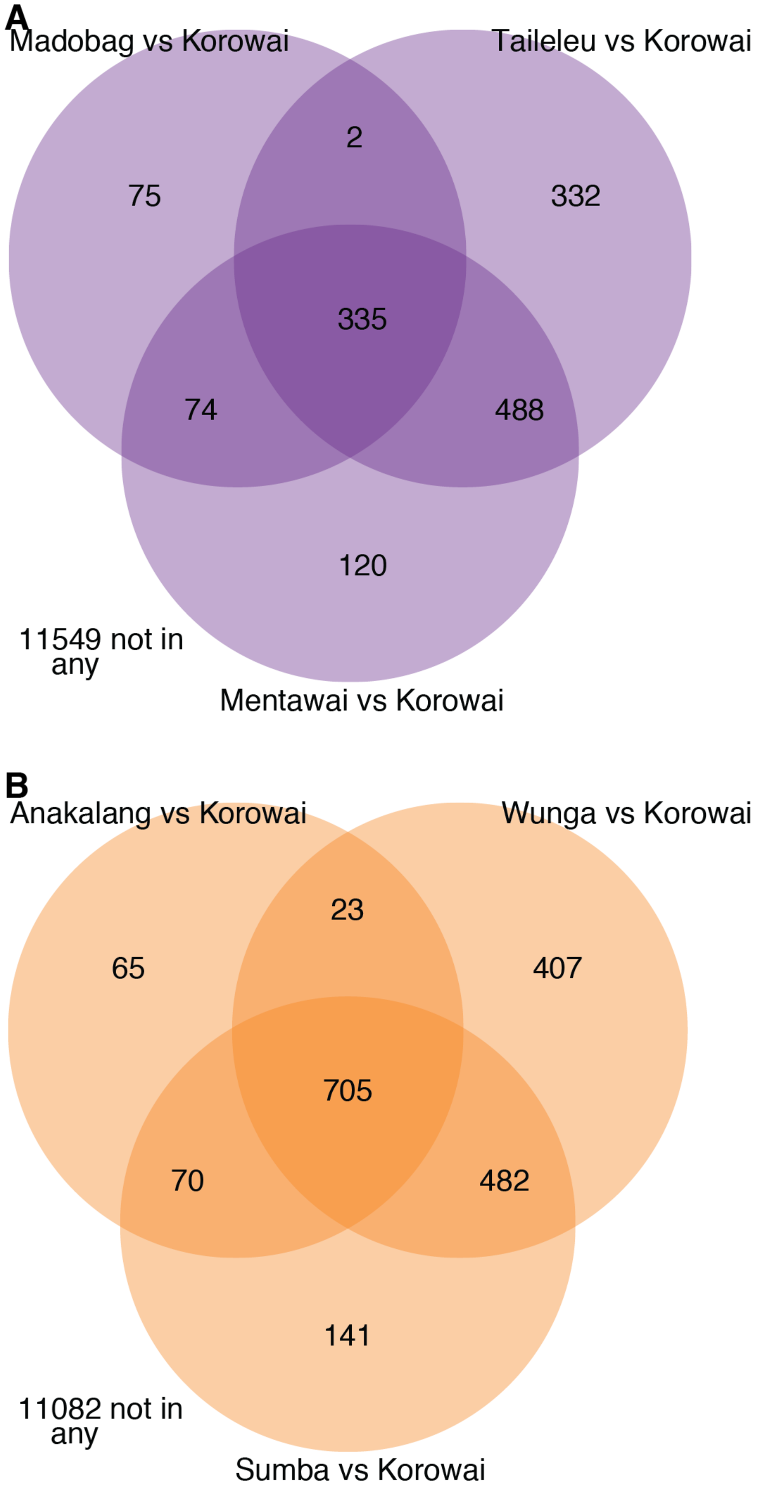


**Supplementary Figure 8:** Sharing of DE signals at the island and village levels in (A) Mentawai vs West Papua comparisons and (B) Sumba vs West Papua comparisons. All comparisons are presented at an FDR threshold of 1% and a $\log_2$ FC threshold of 0.5 or greater.

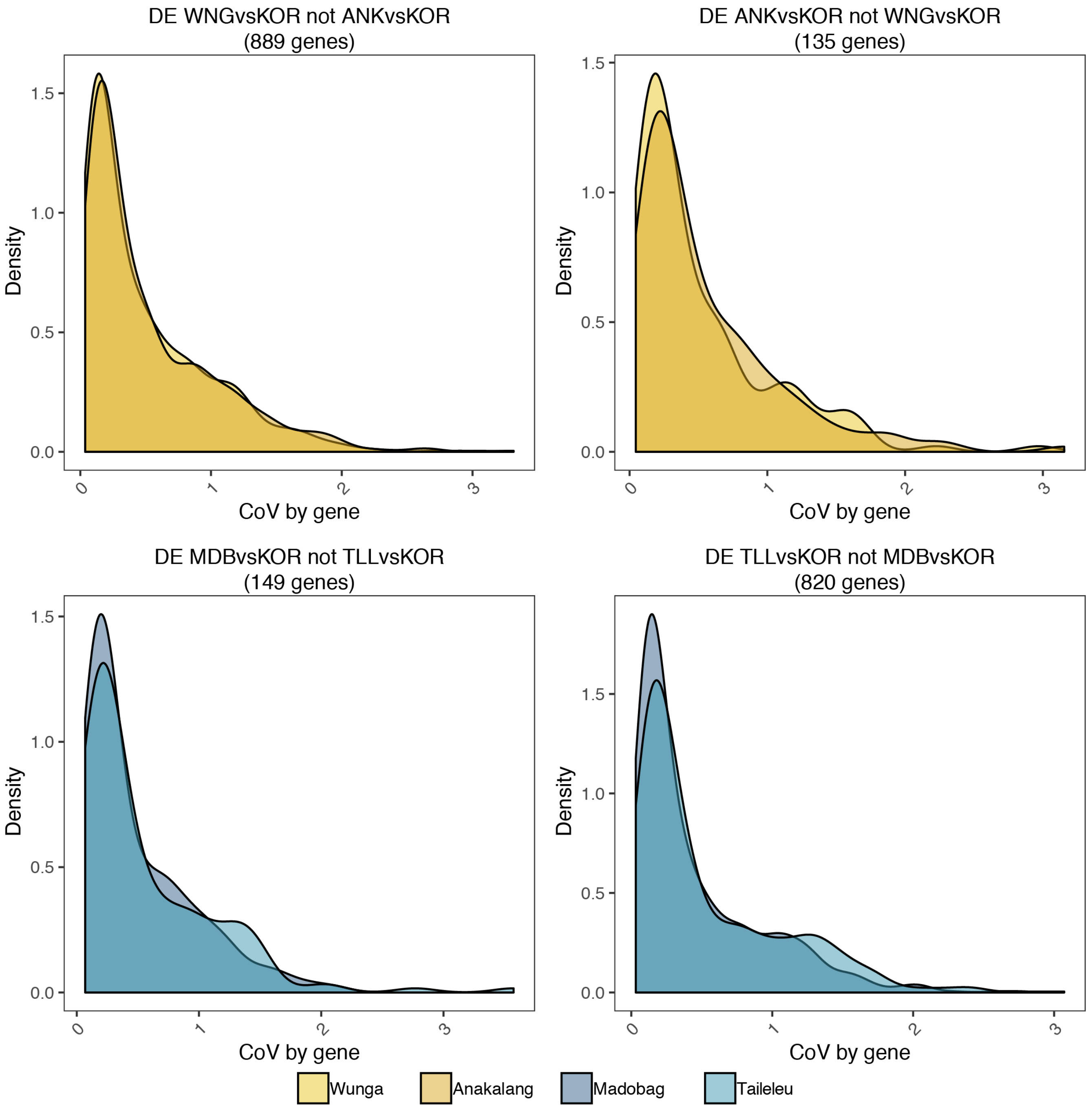


**Supplementary Figure 9:** Distribution of coefficients of variation (CoV) across villages for genes identified as differentially expressed against the Korowai in one village from (top) Sumba or Mentawai (bottom) but not the other.

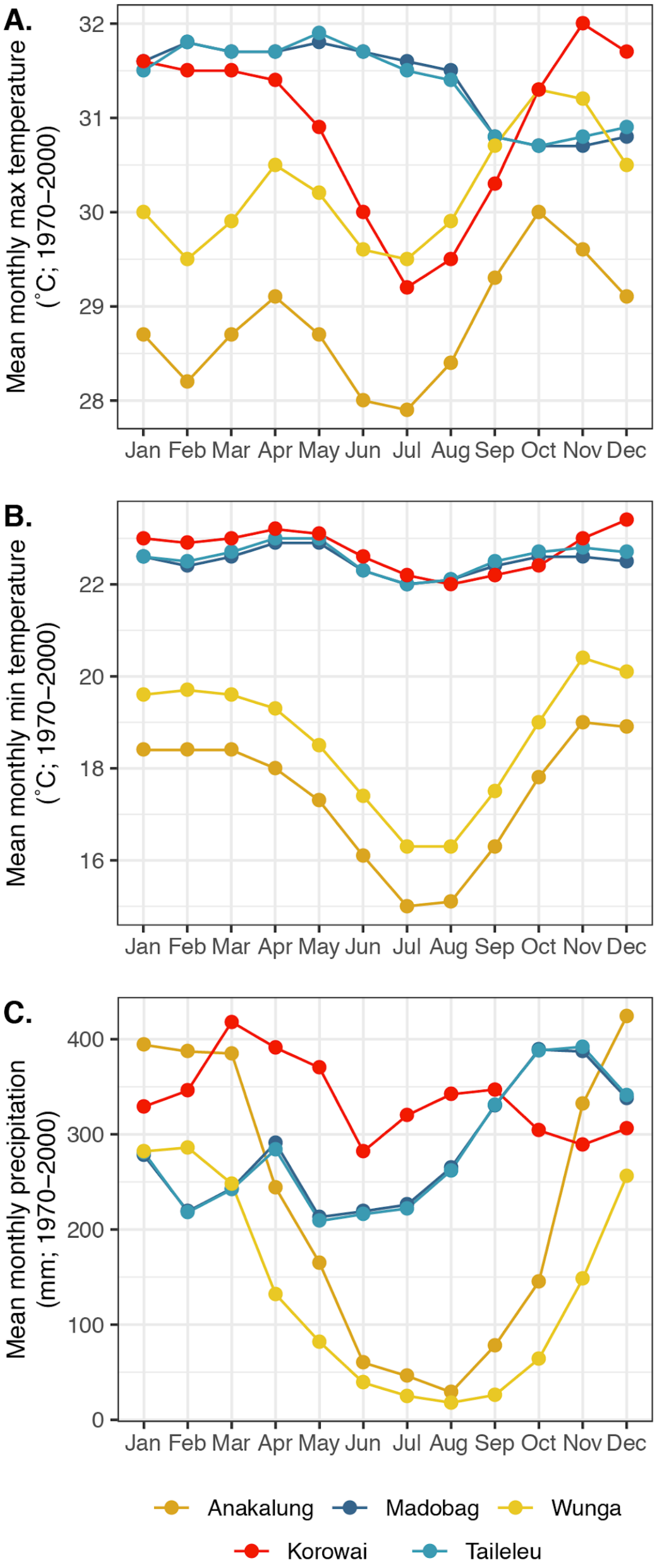


**Supplementary Figure 10:** Monthly climate fluctuations across the five main village sampling sites averaged across 1970 and 2000 retrieved from the WorldClim database. (A) Mean monthly maximum temperature; (B) Mean monthly minimum temperature; (C) Mean monthly precipitation levels.

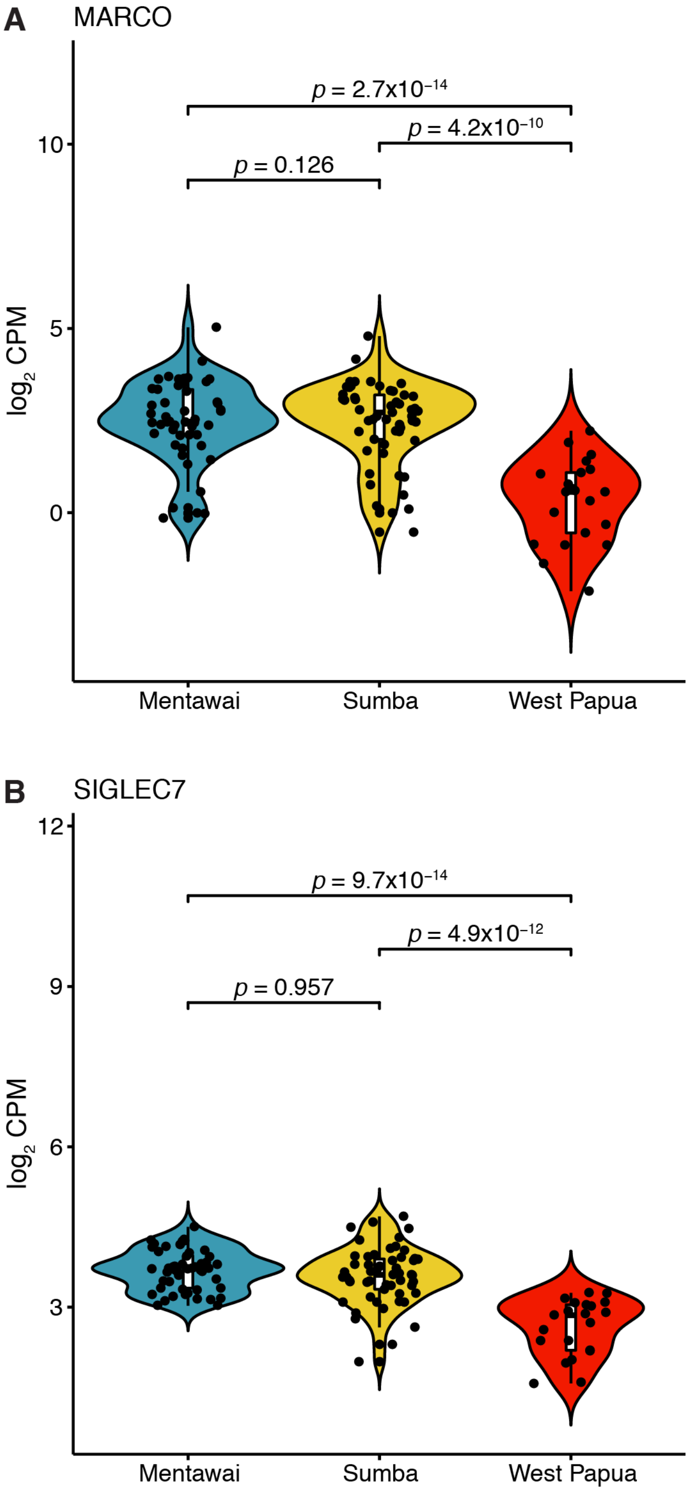


**Supplementary Figure 11:** $\log_2$ CPM values across all samples for (A) *MARCO* and (B) *SIGLEC7*.
